# Differential weighting of information during aloud and silent reading: Evidence from representational similarity analysis of fMRI data

**DOI:** 10.1101/2024.02.18.580744

**Authors:** Lyam M. Bailey, Heath E. Matheson, Jonathon M. Fawcett, Glen E. Bodner, Aaron J. Newman

## Abstract

Single word reading depends on multiple types of information processing: readers must process low-level visual properties of the stimulus, form orthographic and phonological representations of the word, and retrieve semantic content from memory. Reading aloud introduces an additional type of processing wherein readers must execute an appropriate sequence of articulatory movements necessary to produce the word. To date, cognitive and neural differences between aloud and silent reading have mainly been ascribed to articulatory processes. However, it remains unclear whether articulatory information is used to discriminate unique words, at the neural level, during aloud reading. Moreover, very little work has investigated how other types of information processing might differ between the two tasks. The current work used representational similarity analysis (RSA) to interrogate fMRI data collected while participants read single words aloud or silently. RSA was implemented using a whole-brain searchlight procedure to characterize correspondence between neural data and each of five models representing a discrete type of information. Both conditions elicited decodability of visual, orthographic, phonological, and articulatory information, though to different degrees. Compared with reading silently, reading aloud elicited greater decodability of visual, phonological, and articulatory information. By contrast, silent reading elicited greater decodability of orthographic information in right anterior temporal lobe. These results support an adaptive view of reading whereby information is weighted according to its task relevance, in a manner that best suits the reader’s goals.

## Overview

Single word reading is an automatic and effortless process for most literate individuals. From a cognitive perspective, however, word reading may be considered as a series of computations whereby a printed stimulus is mapped to cognitively relevant representations. In order to characterise the neural underpinnings of word reading, it is useful (and, indeed, commonplace in cognitive neuroscience) to decompose this complex process into discrete types of information processing, while recognizing insights from recent language models which reveal complex interactions between different levels of representation (e.g., Caucheteux et al., 2023; Henningsen-Schomers & Pulvermüller, 2022). In this spirit, we consider five broadly-defined types of information that may be extracted from a printed word: visual, orthographic, phonological, semantic, and (in the case of reading aloud) articulatory. While neural processes concerning each type of information have been studied extensively, it remains unclear how, and to what extent they might differ between reading aloud and reading silently.

This question is particularly relevant in the context of embodied and grounded cognition; that is, the perspective that our internal cognitive states are influenced (if not determined) by the manner in which we receive input from and interact with our environment (Matheson & Barsalou, 2018). From this perspective, viewing cognitive processes (such as reading) as being fundamentally experience-dependent is essential for understanding their neural substrates. In many respects, reading aloud and reading silently entail fundamentally different experiential (and, therefore, cognitive) states. First and foremost, reading aloud entails additional motoric / articulatory processes required for speech production. In turn, the physical act of articulation elicits sensory experiences: the sensation of moving one’s mouth and tongue, feeling one’s larynx vibrate, and hearing one’s own voice. Moreover, reading aloud has been shown to confer a reliable memory advantage (MacLeod et al., 2010), suggesting differences in cognitive processing beyond low-level motoric and sensorimotor mechanisms. This *production effect* is often attributed to distinctive sensory experiences brought about through articulation, though other mechanisms have been proposed. For example, there is evidence that words read aloud benefit from more elaborate semantic processing (Fawcett et al., 2022), which may facilitate encoding and retrieval. All of this is to say that differences between aloud and silent reading extend beyond the mere physical act of articulation, and entail extensive cognitive differences. Although some work has investigated this possibility from a cognitive-behavioural perspective, it remains unclear how such differences might manifest at the neural level.

This question may be addressed with neuroimaging. In particular, representational similarity analysis (RSA) is a technique that allows us to decode different types of information (defined by formal hypothesis models) in neural patterns. The purpose of the present study was to investigate the presence of information in neural patterns associated with different types of information present in printed words, by applying RSA to functional magnetic resonance imaging (fMRI) data acquired during aloud and silent reading.

Below we provide a brief overview of the functional neuroanatomy of single word reading, emphasising cognitive and neural processes associated with each type of information described above. This is intended to provide some context for RSA literature on single-word reading and, ultimately, the design of the current study.

### Functional Neuroanatomy Of Single Word Reading

When presented with a printed or written word, an individual must process its low-level visual properties—shape and orientation of the constituent letter strokes, size, colour, etc. At the neural level, these perceptual processes are largely governed by primary and associative visual cortices housed in occipital cortex (Cornelissen et al., 2009; Gramfort et al., 2012; Tarkiainen et al., 1999). Distinct from this low-level perceptual processing is orthographic processing—the recognition of multi-character strings as visual word-forms. Visual word-forms may be described as perceptually invariant mental representations of words, irrespective of size, color, font, or position in the visual field (Warrington & Shallice, 1980). With respect to the neural correlates of orthographic processing, much emphasis has been placed on the visual word form area (VWFA), located in the tempero-occipital portion of the left fusiform gyrus. Neuroimaging work has demonstrated that this area responds preferentially to both real words and orthographically regular pseudowords compared to irregular consonant-string pseudowords (Cohen et al., 2002; Petersen et al., 1990; Polk & Farah, 2002), and also to real words and consonant strings compared to false font or unknown character strings (Baker et al., 2007; Brem et al., 2010; Carreiras et al., 2014; Pleisch et al., 2019). These findings indicate that VWFA is sensitive both to letter strings generally, relative to perceptually similar non-letter visual objects, and also to orthographic regularity. In other words, it appears specialised for detecting visual stimulus properties that adhere to learned orthographic rules (i.e., statistics of written language), thus enabling recognition of familiar visual word forms. Moreover, studies on individuals suffering from pure alexia (selective impairment of the ability to recognise words rapidly and automatically, while identification of individual letters is preserved; McCandliss et al., 2003) often report damage to VWFA (Pflugshaupt et al., 2009; Turkeltaub et al., 2014), suggesting a causal role of this area in orthographic processing.

During reading, orthographic information is associated to an auditory phonological code—that is, a mental representation of the constituent sounds that make up the spoken form(s) of the word they are reading (Leinenger, 2014). The nature of phonological processing in reading has historically been contentious, with opposing connectionist (e.g., Seidenberg & McClelland, 1989) and dual-route (e.g., Coltheart et al., 2001) models describing different mechanisms (see Seidenberg et al., 2022 for discussion). Here we adopt the connectionist principle of weighted spelling-to-sound mapping (Seidenberg, 2005)—which has been integrated into more recent connectionist-dual-process (CDP) models of word reading, and whose purpose is to bridge the gap between the two accounts (e.g. the CDP++ model; Perry et al., 2010). From this perspective, phonological forms are computed based on weighted connections between orthographic units and sounds; the weightings themselves are determined by the consistency of spelling-to-sound mappings within the language in question^1^. Words with inconsistent mappings may be considered to impose greater demands on the spelling-to-sound conversion system; hence, structures that are sensitive to spelling-sound consistency likely play a role in this process. A number of fMRI studies have revealed preferential activation of the left inferior frontal gyrus, and neighbouring structures such as anterior insula and anterior cingulate, when participants see words with inconsistent spelling-to-sound mappings relative to those with consistent mappings, implicating these structures in spelling-to-sound mapping (Bolger et al., 2008; Fiez et al., 1999; Fiez & Petersen, 1998; C.-Y. Lee et al., 2004).

Readers also experience semantic processing — that is, rapid and automatic retrieval of the meaning of the word they are reading. Based on a meta-analysis of 180 fMRI studies (Binder et al., 2009), Binder and Desai (2011) proposed a left-lateralized neurobiological model of semantic processing comprising dorsomedial and inferior prefrontal, inferior parietal, and inferior and ventral temporal cortices. Some of these same areas (particularly the inferior frontal and inferior temporal gyri, and the inferior parietal lobule) have also been identified by other meta-analyses of semantic processing (Rodd et al., 2015; Vigneau et al., 2006).

In the case of reading aloud, an individual must engage additional articulatory (motor) processes, and will experience both proprioceptive feedback from moving one’s mouth and tongue, and acoustic stimulation associated with hearing the sound of one’s own voice. Indeed, a number of fMRI studies have reported that, compared to silent reading, reading aloud is associated with activation of auditory and sensorimotor cortices (Bailey et al., 2021; Dietz et al., 2005; Qu et al., 2022). Bailey et al. attributed their findings to motoric and sensory experiences involved in articulation, while Dietz et al. emphasised increased phonological processing demands in the context of reading aloud. In addition, a meta-analysis by Murphy et al. (2019) identified a single cluster in left STG that responded preferentially to aloud compared to silent reading. Although the authors did not offer an interpretation of this finding, one might take left STG activation to reflect auditory processing while reading aloud, given that STG has been implicated in both basic auditory processing, and speech production specifically (Hickok & Poeppel, 2007; Scott et al., 2000).

### Investigating Aloud And Silent Reading With RSA

Recent years have seen growing popularity of multivariate analysis methods for fMRI, which are all broadly concerned with the distributed patterns of activation elicited by stimuli. One such approach is RSA (Kriegeskorte et al., 2008), which allows us to characterise the informational content of these distributed activation patterns. Rather than making inferences based on contrasts between experimental conditions (as in univariate studies described above), RSA permits direct comparison between neural activation patterns and explicit hypothesis models, which often characterise specific stimulus information. Formally, this method quantifies the representational geometry of stimulus-specific activation patterns—that is, the geometric relationships between stimuli in high-dimensional space—as a *representational dissimilarity matrix* (RDM). This neural RDM may then be compared to one or more researcher-defined RDMs (hypothesis models) derived from quantitative measures of a given stimulus property. Within this framework, we can decode (i.e., detect) specific types of stimulus information (visual, orthographic, semantic, etc.) in a given patch of cortex, by demonstrating statistical dependence between a neural RDM (extracted from patterns in that patch) and a relevant hypothesis model (Kriegeskorte & Kievit, 2013).

A number of fMRI-RSA studies have investigated decodability of different types of information during single word reading. To our knowledge, only one study has examined low-level visual information in this context—Fischer-Baum et al. (2017) reported visual decodability in right posterior ventral visual cortex, consistent with RSA studies using images as stimuli which reliably find visual information in posterior visual areas (e.g., Devereux et al., 2013; Kriegeskorte et al., 2008; Proklova et al., 2016). Meanwhile, orthographic information has been decoded from activity patterns in the left temporal lobe, including the tempero-occipital portion of the fusiform gyrus (which notably houses putative VWFA; Fischer-Baum et al., 2017; Qu et al., 2022), as well as more anterior portions of ventral and lateral temporal lobe (Graves et al., 2023; Staples & Graves, 2020; Zhao et al., 2017). Beyond the temporal lobe, orthographic information has also been detected in the inferior (Fischer-Baum et al., 2017) and superior (Graves et al., 2023) parietal lobules, while a pair of studies reporting whole-brain analyses (Graves et al., 2023; Staples & Graves, 2020) detected orthographic information in left prefrontal and medial occipital cortices.

Phonological information has been decoded across a similarly broad number of areas (in some cases overlapping with orthographic information; Graves et al., 2023); in left ventral and lateral temporal cortex (Fischer-Baum et al., 2017; Graves et al., 2023; Qu et al., 2022; Staples & Graves, 2020; Zhao et al., 2017), left prefrontal and inferior parietal cortices (Fischer-Baum et al., 2018; Graves et al., 2023; Li et al., 2022), as well as middle and superior lateral temporal cortex (Graves et al., 2023; Li et al., 2022; Staples & Graves, 2020).

Semantic information has been decoded in ventral tempero-occipital cortices (Fischer-Baum et al., 2018; Wang et al., 2018), the left inferior parietal lobule (Fischer-Baum et al., 2018; Graves et al., 2023), left middle frontal and superior temporal gyri, and medial and lateral occipital cortex bilaterally (Graves et al., 2023). To our knowledge, only one study has investigated articulatory information in the context of reading: Zhang et al. (2020) had participants either read aloud, mouth, or imagine reading aloud simple consonant-vowel syllables, and reported that articulatory and acoustic information was decodable across a range of frontal, temporal, and parietal ROIs previously implicated in speech production and sensation.

Taken together, the studies described above indicate that each type of information that we consider relevant to word reading may be decoded in multiple brain areas during single word reading. Most of these studies have focussed on aloud or silent reading tasks independently; a critical question, however, is whether the decodability of various types of information might *differ* between these two tasks. This question is particularly relevant to research on neural correlates of the production effect, which is defined as a contrast between aloud and silent reading. Moreover, from an embodied and grounded cognition perspective, this question is important for assessing the degree to which the neural correlates of reading are experience-dependent.

Decoding-based differences between aloud and silent reading would be consistent with prior RSA work showing that semantic decodability in particular is often task- or experience-dependent (e.g., when performing tasks that emphasise different semantic features of presented stimuli; Meersmans et al., 2022; Nastase et al., 2017; Wang et al., 2018). It is possible that decodability of other types of information is similarly variable when comparing aloud to silent reading. Indeed, there are many reasons to expect such differences, outlined as follows. First and perhaps most obviously, one might expect greater dependence on (and therefore increased decodability of) articulatory information during aloud versus silent reading, because speaking a word necessarily requires participants to execute an appropriate sequence of articulatory movements. In this context, articulatory information should be decodable in primary motor and/or premotor cortices (i.e., precentral gyrus and supplementary motor area). Indeed, previous fMRI work has shown that vocalising different phonemes elicits discriminable responses in the precentral gyrus (Pulvermüller et al., 2006); this area may therefore have the capacity to represent whole words as linear combinations of articulatory features that are required to produce the constituent sounds.

Neural mechanisms distinguishing aloud and silent reading likely extend beyond the motoric components of articulation. Cognitive work surrounding the production effect—whereby words read aloud are more readily remembered compared to words read silently (MacLeod et al., 2010)—has emphasised the role of distinctive sensorimotor experiences during articulation. These include the sensation of moving one’s mouth and tongue, feeling one’s larynx vibrate, and experiencing auditory feedback from hearing one’s own voice. At a higher cognitive level, reading aloud also entails planning and monitoring of speech output, and possibly sensory attenuation of efferent signals (corollary discharge) (Khalilian-Gourtani et al., 2022). At the neural level, we might expect such distinctiveness to be manifest as changes in decodability of articulatory and/or phonological information, perhaps in higher-level (i.e., non-sensorimotor) areas associated with speech planning and monitoring, or episodic encoding. Broadly speaking, the former processes have been linked to medial and lateral prefrontal cortices (Bourguignon, 2014; Hertrich et al., 2021), while episodic encoding is thought to depend largely on frontoparietal and medial temporal areas (e.g., H. Kim, 2011).

There is also evidence that reading aloud enhances semantic processing (Fawcett et al., 2022). We might therefore expect to see higher semantic decodability during aloud versus silent reading. It has also been suggested that people allocate more attention to words that they read aloud (Fawcett, 2013; Mama et al., 2018; Ozubko et al., 2012). While attention is not clearly defined in this context (particularly in terms of its relationship to specific types of information), we might broadly consider this term to reflect changes in one’s cognitive state whereby words read aloud are assigned greater weighting (i.e., they are prioritised) during perceptual and goal-directed cognitive operations, compared to words read silently. This interpretation is consistent with prior conceptualizations of attention as the allocation of (limited) cognitive resources to a stimulus (P. A. MacDonald & MacLeod, 1998; Mama et al., 2018). Upon seeing a cue to read an upcoming word aloud, participants may experience an overall increased level of arousal as they prepare to map upcoming visual/orthographic information onto an appropriate vocal response. Increased arousal may lead to greater cognitive “investment” in processing multiple types of information (and therefore increased decodability of each), recognising that successful word production depends on accurate mapping of low-level visual information to orthography, phonology, and articulatory features.

The predictions outlined above have received partial support from two studies examining task-dependent changes in decodability during single word reading. Qu et al. (2022) reported that, relative to silent reading, aloud reading elicited greater decodability of orthographic and phonological information in the left anterior fusiform gyrus. Moreover, Zhang et al. (2020) presented participants with consonant-vowel syllables and reported that reading aloud, mouthing, and imagined reading aloud elicited differential correlations with an articulatory model in a number of areas implicated in speech production and sensation. Notably, articulatory information was decodable in left angular and superior temporal gyri when reading aloud, but not in the other two conditions. These two studies provide proof-of-principle that different reading tasks may modulate decodability of phonological, orthographic, and articulatory information. However, a number of questions still remain. Qu et al.’s (2022) analysis was confined to the fusiform gyrus; therefore, it is unclear how phonological and orthographic information (or, indeed, other types of information) might be differentially represented in other brain areas. Moreover, Zhang et al. (2020) did not explicitly compare aloud reading with silent reading (rather, these authors examined three production tasks with variable degrees of enactment), and so it remains unclear how articulatory decodability might differ between the former conditions.

### The Current Study

In the above section, we showed that different kinds of information—visual, orthographic, phonological, semantic, and articulatory—may be decoded during single word reading. There is some evidence for differential decodability of phonological and orthographic information between aloud and silent reading in the fusiform gyrus (Qu et al., 2022), but whether similar effects are present in other areas, or for other types of information, remains unclear. As such, the goal of the current work was to establish how reading words aloud versus silently affects the decodability of visual, orthographic, phonological, semantic, and articulatory information throughout the whole brain. We conducted an fMRI experiment in which participants saw visually presented words (each repeated multiple times throughout the experiment) and were instructed to read each word either aloud or silently. We used a searchlight procedure to generate a neural dissimilarity matrix centred on each voxel throughout the brain^2^; we then compared the neural data from each searchlight to hypothesis models representing each of the five types of information discussed above.

Given that articulatory information is essential for speech production, we predicted that, relative to silent reading, reading words aloud would result in increased articulatory decodability in primary and associative motor areas. Articulatory (and possibly phonological) information may also be present in frontoparietal and superior temporal areas associated with speech planning and monitoring. These types of information may also be present in medial temporal areas associated with episodic encoding, consistent with accounts of the production effect which emphasise sensorimotor experiences during encoding of words spoken aloud. From the view that reading aloud enhances semantic processing, we predicted increased semantic decodability relative to silent reading. While univariate studies have implicated multiple areas in semantic processing, changes in semantic decodability seems most likely in ventral tempero-occipital cortex, as this area has been implicated by multiple RSA studies. Finally, increased attention to aloud words may lead to increases in decodability of multiple types of information. Given that ventral tempero-occipital cortex (particularly the left fusiform gyrus) has attracted much attention concerning visual, orthographic, and phonological decodability, it seems plausible that such attentional effects would be present in this area. All of these predictions would be evidenced by higher correlations between the relevant hypothesis model(s) and neural patterns elicited by aloud, compared to silent reading.

## Methods

### Subjects

Data was collected from 30 participants, aged 18-40 (*M* = 21.43, *SD* = 4.57), 21 female, three left-handed. We chose to include left-handed participants in light of growing calls for more inclusive research practices, particularly with respect to handedness (Bailey et al., 2019; Willems et al., 2014) (Bailey et al., 2019; Willems et al., 2014). We (Bailey et al., 2019) have previously argued that any deleterious effects of including left-handers in a sample of 20-30 participants, even in the context of a language study, will be negligible. All participants reported normal or corrected-to-normal vision, proficiency in English, no history of neurological illness or trauma, and no contraindications to MRI scanning. Handedness information was obtained using the Edinburgh Handedness Inventory (Oldfield, 1971). Participants were recruited through on-campus advertising at Dalhousie University, and received $30 CAD reimbursement and a digital image of their brain. All data collection took place at the IWK (Izaak Walton Killam) Health Centre in Halifax, NS. All procedures were approved by the IWK Research Ethics Board. Participants provided informed consent according to the Declaration of Helsinki.

Four participants were excluded from data analysis: one reported difficulty reading visually presented words while in the scanner; one reported that they did not follow task instructions properly (they silently mouthed words in the aloud condition, instead of vocalising the words as instructed); one disclosed that they did not fit inclusion criteria (neurological normality) after having completed the study; one withdrew prior to completing all parts of the experiment. Therefore, data from 26 participants (18 female, two left-handed) were included in our analyses.

### Stimuli And Apparatus

Participants viewed all stimuli while lying supine in the MRI scanner; stimuli were presented using the VisualSystem *HD* stimulus presentation system (Nordic Neuro Lab, Bergen, Norway). Stimuli were presented on an LCD screen positioned behind the scanner and viewed by participants via an angled mirror fixed to the MR head coil. During the experiment, participants made button-press responses using MR-compatible ResponseGrip handles (Nordic Neuro Lab, Bergen, Norway), one in each hand. Stimuli were presented using PsychoPy 2020.2.1 (Peirce et al., 2019). We selected a subset of 30 nouns from Bailey et al. (2021) which, in turn, were sourced from MacDonald and MacLeod (1998). Words were 6 to 10 characters in length, each with a frequency greater than 30 per million (Thorndike & Lorge, 1944). Our full word list is presented in Supplementary Table 1.

All words were presented at the center of the screen in white lowercase Arial font against a dark grey background (RGB: 128, 128, 128). Response instructions for each word (see Procedure) were grayscale icons presented at the start of each trial in the centre of the screen—participants were instructed to speak the word aloud if they saw a mouth icon, or to read the word silently if they saw an eye icon. For our active baseline task (see Procedure), individual numbers 1-9 were presented in white Arial font at the centre of the screen against the same grey background as the words.

#### Hypothesis Models

We constructed a set of hypothesis models used for RSA (see 2.6) reflecting visual, orthographic, phonological, semantic, and articulatory properties of the words presented in our experiment. Unique hypothesis models were generated for each participant and condition separately (because words were randomly allocated to conditions for each participant, see 2.3), in the Python environment using custom scripting and publicly available Python packages. Each hypothesis model comprised a 15 x 15 representational dissimilarity matrix (RDM) containing zeros along the diagonal; off-diagonal cells contained pairwise dissimilarity values corresponding to all possible pairs of words within that condition, and matrices were symmetrical about the diagonal (Kriegeskorte et al., 2008). To generate each model, we computed dissimilarity values according to measures that were theoretically relevant to the respective type of information / stimulus property being examined. Each of these measures is briefly outlined below, with more details provided in Supplementary Materials. Hypothesis models from a representative subject and condition are shown in Figure 1.

**Figure 1.**
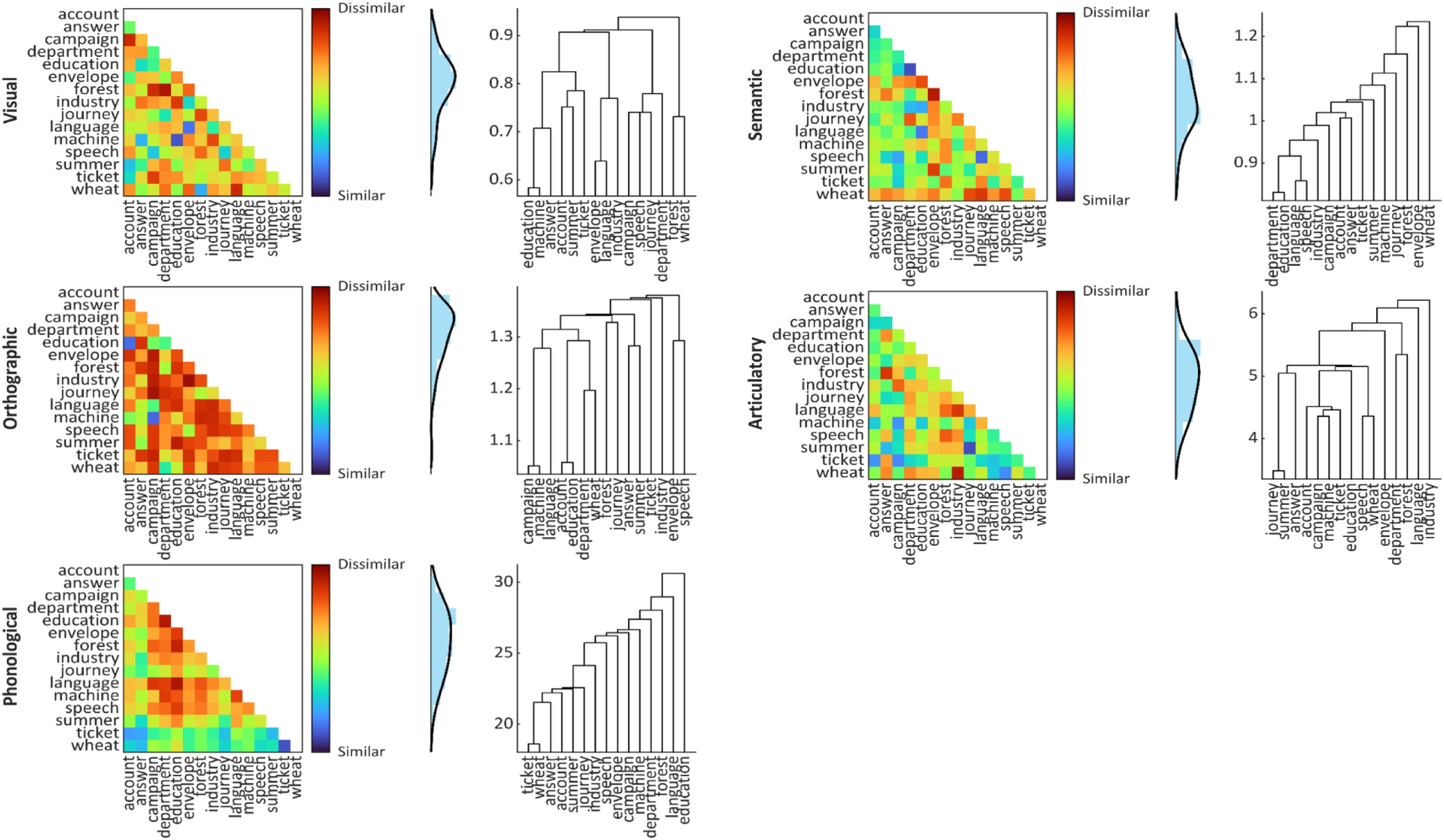
Hypothesis models (i.e., representational dissimilarity matrices; RDMs) from a representative subject and condition. Values shown in heatmaps and histograms have been *z*-scored to reflect the data entered as regressors in each GLM.

We computed visual dissimilarity as the correlation distance (1 - Pearson correlation) between vectorized binary silhouette images of each word as it was presented in the experiment (as in Kriegeskorte et al., 2008). This procedure was implemented using the Pillow package (Umesh, 2012). We computed orthographic dissimilarity as the correlation distance between unconstrained open bigram feature vectors (similar to Fischer-Baum et al., 2017), using the wordkit package (Tulkens et al., 2018).

We computed phonological dissimilarity using the measure described by Bartelds (2020)—in brief, an audio sample of each word is decomposed into segments, each of which is represented by a set of Mel-frequency cepstral coefficient (MFCC) features. The dissimilarity between a pair of words is described by Euclidean distances between their constituent sets of features. This measure is therefore best described as one of acoustic dissimilarity. While we are unaware of any studies using this measure as input for RSA, this measure has been shown to accurately predict human judgments of the native-likeness of North American English pronunciations (Bartelds et al., 2020); recognising that native-likeness is a common benchmark for quantifying speech-based similarity (e.g., Fontan et al., 2016). Therefore, we feel that this measure should adequately capture auditory phonological codes discussed in the Introduction. We computed phonological dissimilarity using Python code provided by Bartelds (2020), based on audio samples for each of the words in our stimulus list. To this end we collected voice recordings from seven volunteers (of similar demographic characteristics to our main study sample), as well as four AI-generated voices (simulating Canadian and American speech profiles) provided by the online text-to-speech platform *Vecticon* (https://vecticon.co)^3^. Pairwise phonological distances between words were computed using speech samples from each speaker independently (i.e., we only compared words from the same volunteer / AI voice); these distances were then averaged across speakers to generate a single representative dissimilarity value for every pair of words. Further details are provided in Supplementary Materials.

For semantic models, we computed cosine distances (1 - cosine similarity) between word2vec representations of each word’s semantic content (consistent with previous work; e.g., Carota et al., 2021; Graves et al., 2023; Tong et al., 2022; Wang et al., 2018). Word2vec representations were acquired from the publicly available glove-wiki-gigaword-300 model (https://github.com/RaRe-Technologies/gensim-data) using the gensim package (Řehůřek & Sojka, 2010).

Finally, we computed articulatory dissimilarity as the feature-weighted phonological edit distance (Fontan et al., 2016) between words, normalized for word length (Beijering et al., 2008; Schepens et al., 2012) using the PhonologicalCorpusTools package (Hall et al., 2019). While this is ostensibly a measure of phonological dissimilarity, it depends on articulatory features necessary to produce each word (e.g., place of articulation, movements of the tongue, teeth, and lips, etc.). As such, we feel that it provides a suitable means for modelling articulatory information. Moreover, our analyses required that our five measures be independent from one another (i.e., minimally correlated); as such, we were motivated to select measures of phonological and articulatory dissimilarity that were as different as possible from one another.

To ensure that our five measures were independent from one another, we computed pairwise correlations between (vectorised) exemplar models. Exemplar models were RDMs containing all words in our stimulus set (rather than condition-specific RDMs used in the actual analyses), and were computed for each measure. We reasoned that, because stimulus allocation to each condition was randomised for each participant, comparing exemplar models (containing all possible stimulus combinations) was the best way to approximate potential correlations between models used in our analysis. Correlations between our five exemplar models are displayed in Table 1. Accompanying parenthetical values in Table 1 show Bayes factors BF_10_ (outlined in the Group-Level Analysis section below) for each correlation, computed using the bayesFactor package (Krekelberg, 2022) in the MATLAB environment, using default JZS priors (Rouder et al., 2009). These comparisons revealed strong evidence for a negative correlation between the phonological and semantic models (*r* = −0.23, BF_10_ > 5,000) and even stronger evidence for a positive correlation between the phonological and articulatory models (*r* = 0.29, BF_10_ > 5,000,000); all other comparisons revealed equivocal evidence (BF_10_ < 2.0). The positive correlation between phonology and articulation is to be expected because, from a computational perspective, the constituent sounds of a word are intrinsically related to the articulatory movements necessary to produce those sounds. Why phonology and semantics should be negatively correlated is less clear. Nevertheless, the actual magnitudes of these correlations are low, and both are within the range of equivalent comparisons in previous RSA work (e.g., Staples & Graves, 2020 report inter-model correlations between *r* = 0.52 and 0.63). Moreover, the GLMs used in our searchlight analyses (see Searchlight Analyses section) entailed each model as a regressor; therefore, any overlapping effects between correlated models ought to be partialed out.

**Table 1.** Correlation matrix for the five dissimilarity measures used in this study. Values are correlation coefficients for each pair of measures; values in parentheses are Bayes factors indicating strength of evidence of a true (anti-)correlation.

|  | Visual | Orthographic | Phonological | Semantic | Articulatory |
| --- | --- | --- | --- | --- | --- |
| Visual |  | -0.04<br>(0.05) | 0.11<br>(0.44) | 0.03<br>(0.05) | 0.0005<br>(0.04) |
| Orthographic |  |  | -0.06<br>(0.08) | 0.11<br>(0.45) | 0.13<br>(1.93) |
| Phonological |  |  |  | -0.23<br>(5.46e3) | 0.29<br>(5.71e6) |
| Semantic |  |  |  |  | 0.01<br>(0.04) |
| Articulatory |  |  |  |  |  |

We also considered the potential impact of other, extraneous stimulus properties. Our stimulus list was not controlled in terms of morphological complexity, concreteness, imageability, or grapheme-to-phoneme consistency. In addition, while all of our stimuli were nouns, in some cases their syntactic category (verb or noun) was ambiguous (e.g., *debate, judge, record*). These properties no doubt affect neural processing, but are beyond the scope of this manuscript. We reason that, since stimuli were randomly allocated to conditions, these extraneous properties were unlikely to drive observed differences between aloud and silent reading. Even so, it is possible that such properties might confound our analyses, particularly if they are highly correlated with our properties of interest (visual / orthographic / phonological etc.)—high collinearity between these extraneous properties and our hypothesis models would obfuscate interpretation of our results.

To rule out such collinearity, we constructed 30 x 30 RDMs for each of the following properties: imageability, morphological consistency, concreteness, grapheme-to-phoneme consistency, and syntactic category. Details of each dissimilarity measure is provided in Supplementary Materials. These RDMs were then compared to our exemplar models—all correlations are shown in S Table 2. These comparisons indicated that our phonological measure was positively correlated with grapheme-to-phoneme consistency (*r* = 0.22, BF_10_ = 2072.86), imageability (*r* = 0.21, BF_10_ = 7.35), and morphological complexity (*r* = 0.14, BF_10_ = 3.54), while our semantic measure was positivity correlated with concreteness (*r* = 0.15, BF_10_ = 6.03), and negatively correlated with grapheme-to-phoneme consistency (*r* = −0.14, BF_10_ = 3.38). As before, these correlations are small, and so unlikely to seriously undermine interpretation of our results. All other comparisons revealed equivocal evidence (BF_10_ < 2.0).

**Table 2.**
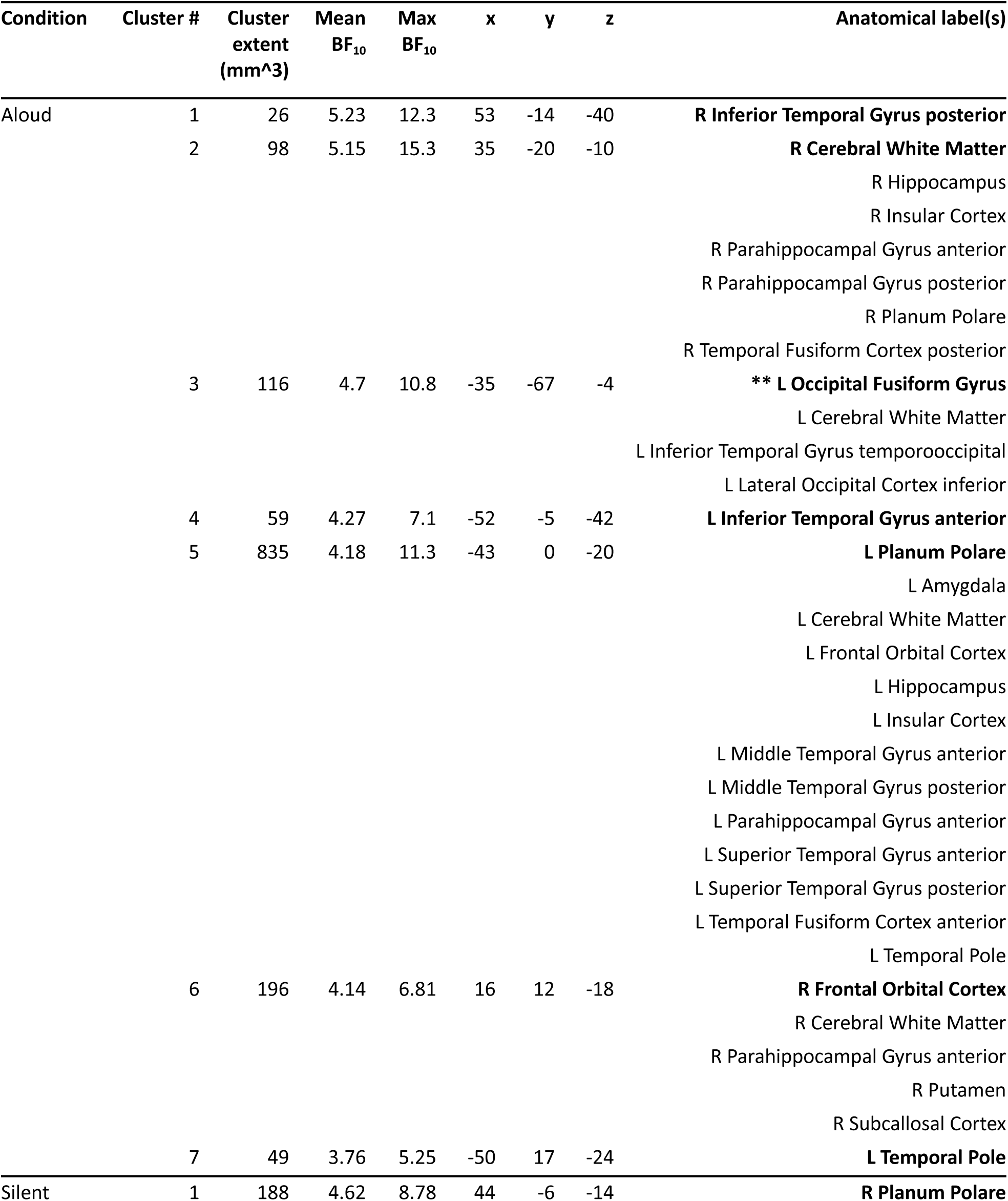
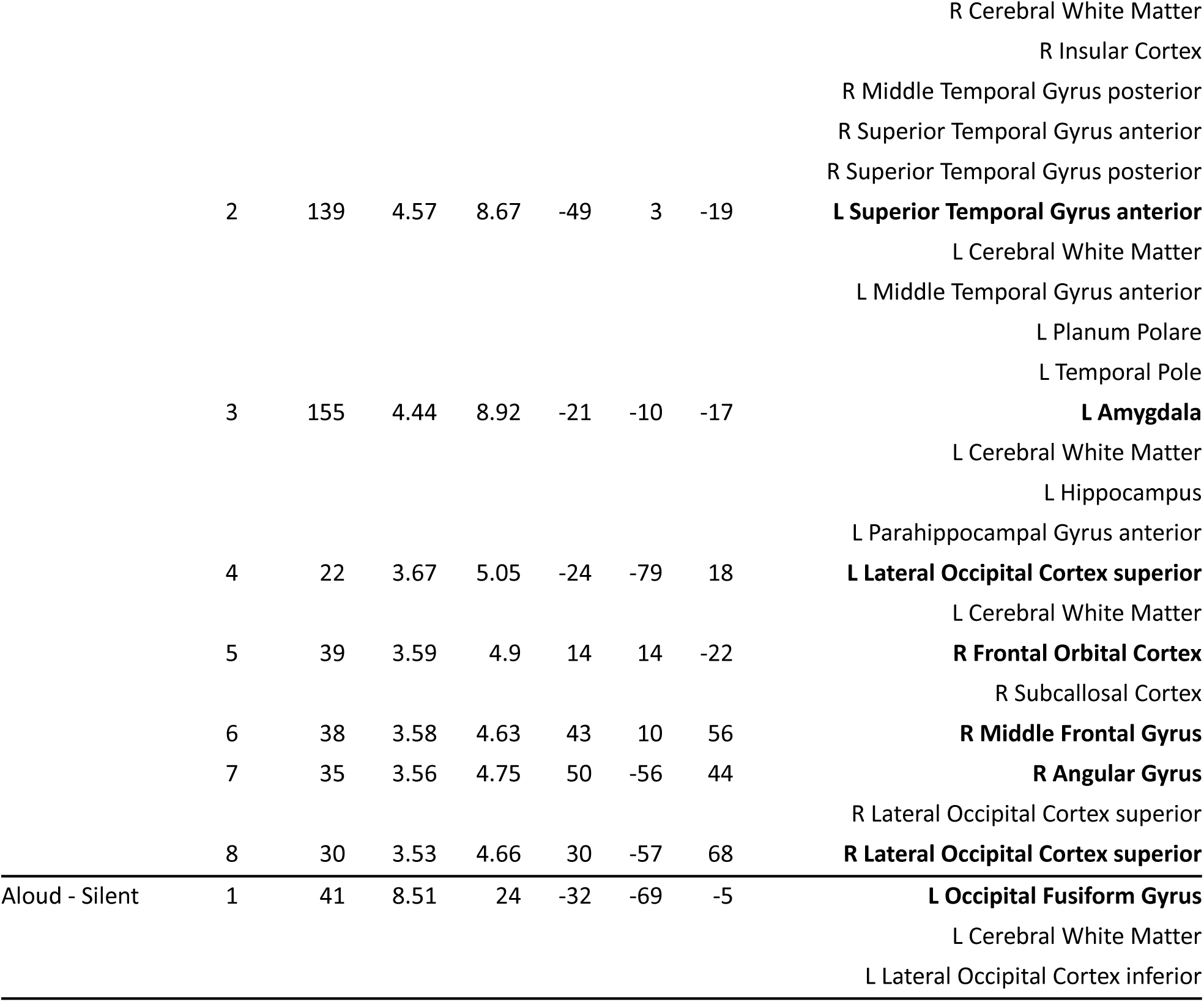
RSA results table detailing clusters for visual information (i.e., elicited by the visual hypothesis model). The anatomical labels column shows anatomical structures encompassed by each cluster; entries in boldface indicate the structure in which the cluster’s center of mass (COM) was situated.

### Procedure

After providing informed consent and completing MRI safety screening and the Edinburgh Handedness Inventory, participants completed a shortened version of the experiment described below (using a different stimulus list) on a laptop computer to familiarise them with the task. Participants were also asked to confirm (verbally) that they understood the task requirements before entering the scanner. Once participants were positioned in the MRI scanner, a brief scout scan determined head position. Following this, participants completed three functional scans as part of another study, followed by a structural scan. After the structural scan, participants completed four three-minute functional scans that comprised the data for the present study.

During each three-minute scan, participants performed a word reading task, in which the 30 words (S Table 3.1) were randomly allocated to one of two conditions (aloud or silent; 15 words in each condition) for each participant. Word-condition mappings were randomised between participants. Each trial of the word reading task began with a fixation cross (“+”) presented for 500 ms to alert the participant that the trial was starting, followed by a cue presented for 1000 ms instructing participants to read the upcoming word either aloud (if they saw a mouth icon) or silently (if they saw an eye icon). Following the cue, a word from the corresponding condition was presented for 2500 ms. Each word appeared once in each functional run, and each of the four runs used a different random order of presentation.

**Table 3.**
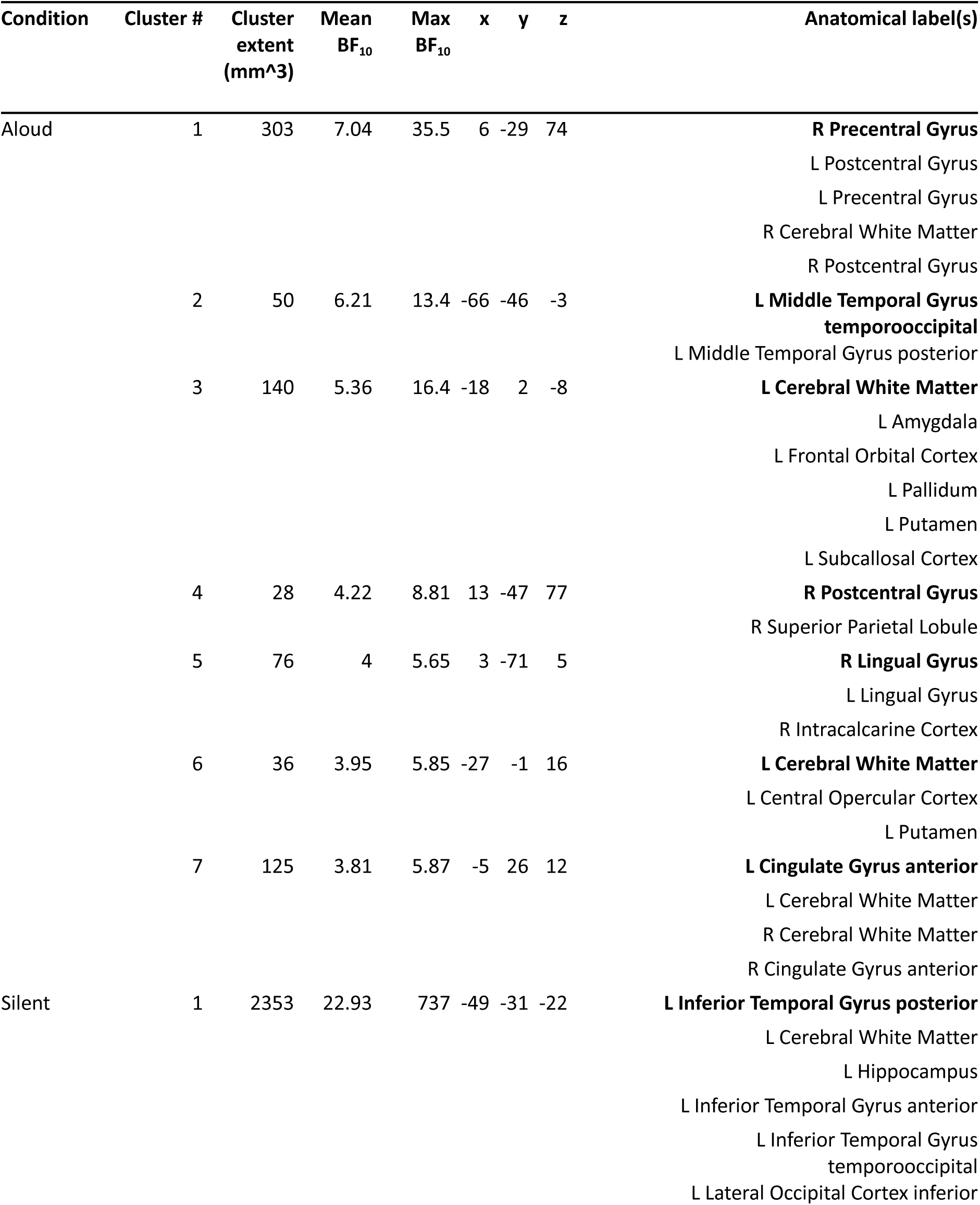
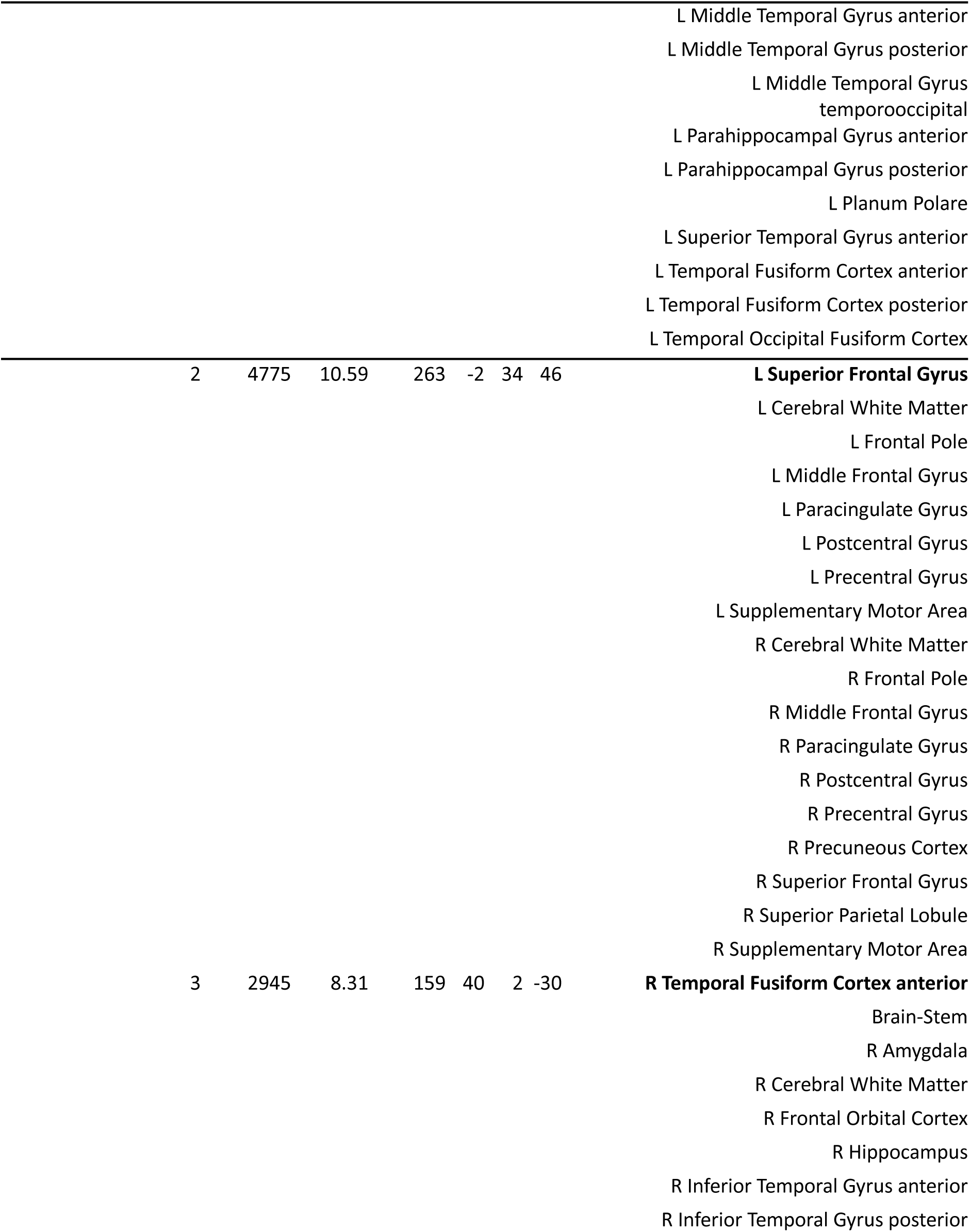
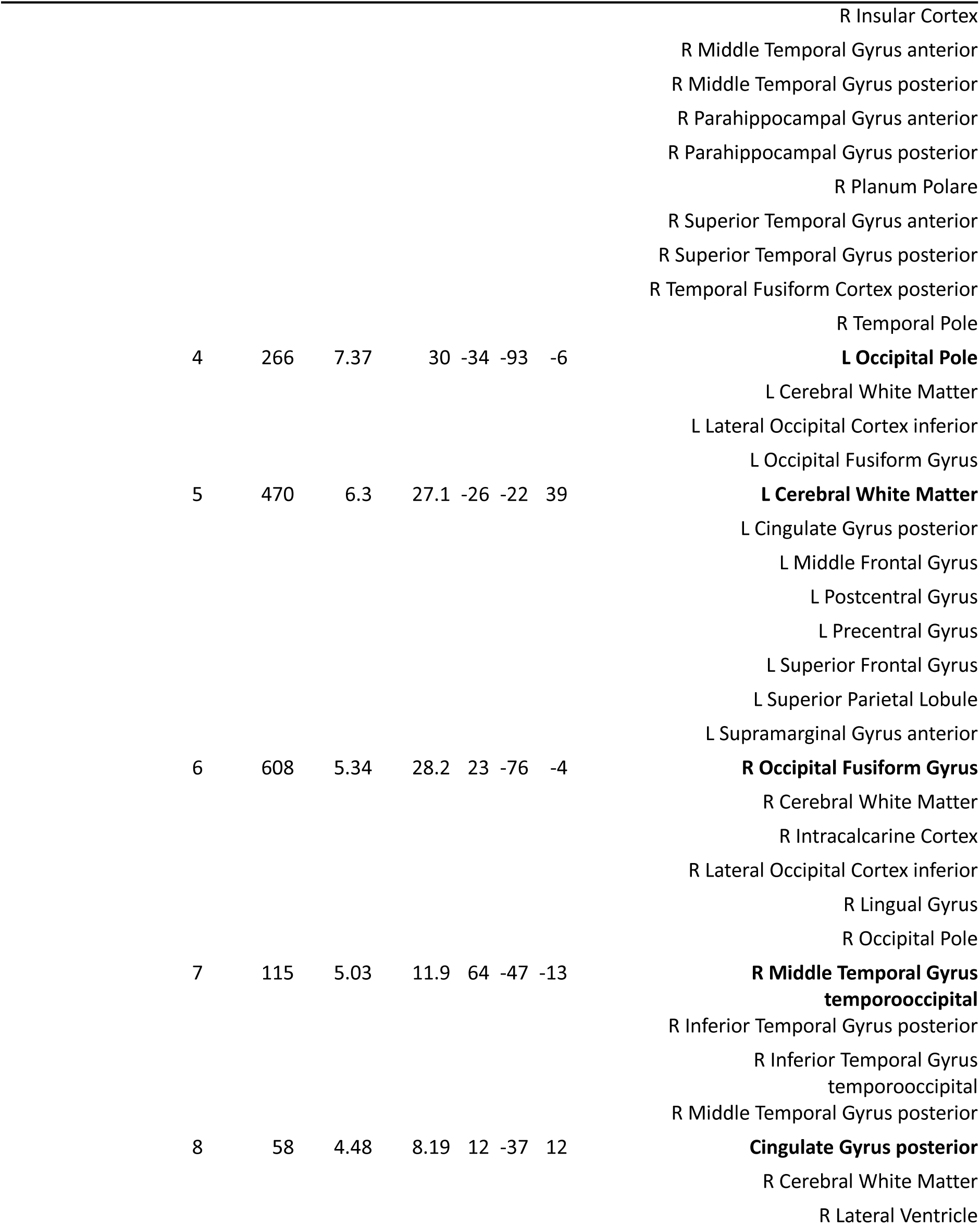
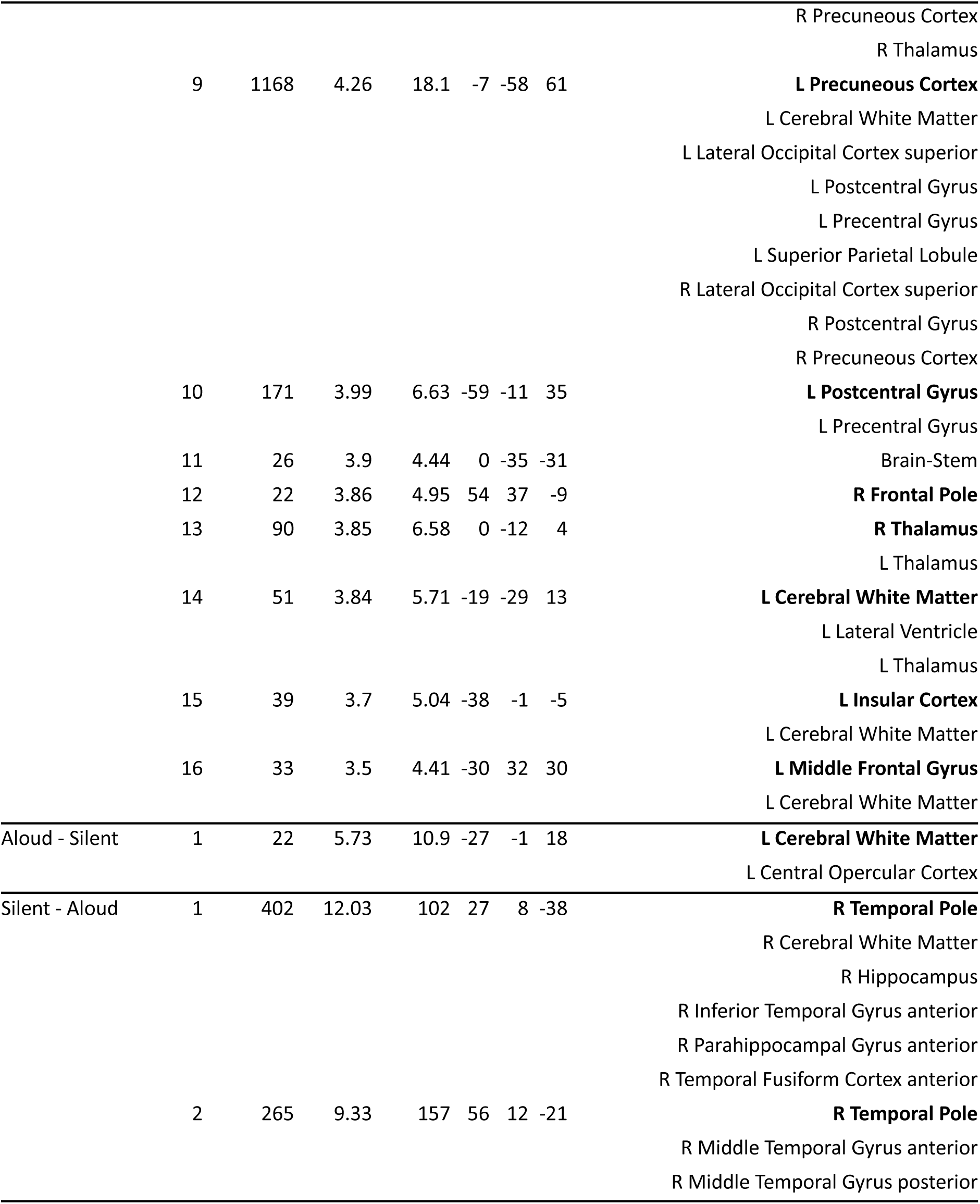
RSA results table detailing clusters for orthographic information (i.e., elicited by the orthographic hypothesis model). The anatomical labels column shows anatomical structures encompassed by each cluster; entries in boldface indicate the structure in which the cluster’s center of mass (COM) was situated.

On every trial, following word presentation, participants completed a short active baseline task: a randomly generated number between 1 and 9 was presented for 2000 ms, and participants were instructed to decide whether the number was odd or even, and to make an appropriate button-press response. During this baseline period, response mapping cues were presented as a reminder in the top corners of the screen (left-hand button press for odd, right-hand for even). Response mappings were the same for all participants. This active baseline served two purposes: (i) to lengthen SOA of each trial to 6 seconds (we did not add temporal jitter or null events—the SOA was always exactly 6 seconds), and therefore ensure reliable item-level estimation (Zeithamova et al., 2017), while (ii) ensuring that participants did not mind-wander between trials. We selected a simple odd-even judgement task because prior work has shown this task to be an effective alternative to the traditional resting baseline in fMRI (Stark & Squire, 2001). A schematic of our trial structure is shown in Figure 2. Each functional run included 30 trials.

**Figure 2.**
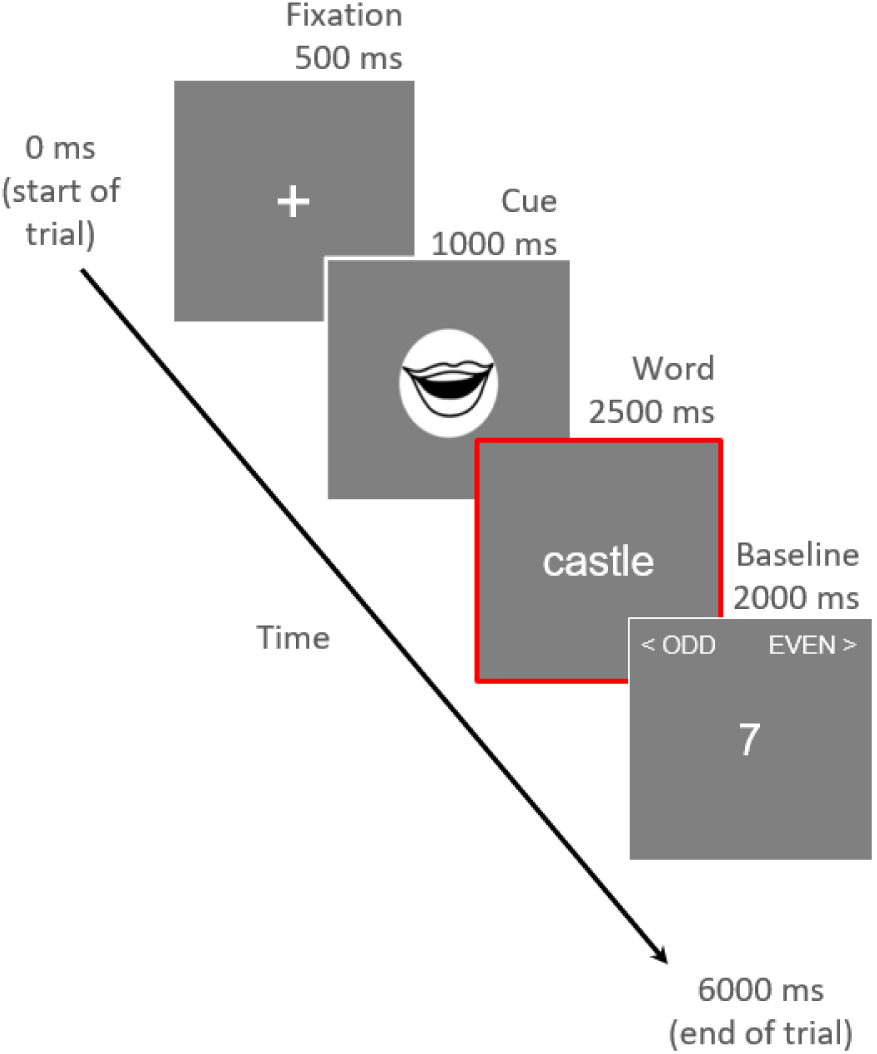
Schematic of an example trial. The red outline indicates the period modelled by the time series regressor for each trial (i.e., the temporal window from which activation patterns were estimated).

### MRI Data Acquisition

MRI data was acquired on a 1.5 Tesla GE MRI system (GE Medical Systems, Waukesha, WI) equipped with a 19 channel head coil. Each participant completed a structural scan followed by four functional scans. As noted above, participants also completed three additional functional scans at the start of their session, but those functional scans are beyond the scope of the current work. For the structural scan, a T1-weighted anatomical image was obtained using a magnetization-prepared rapid acquisition gradient echo (MPRAGE) sequence, inversion time (TI) = 1134 ms, flip angle = 8°, number of excitations (NEX) = 2, field of view (FOV) = 224 mm, matrix = 224 x 224, resulting in an in-plane voxel resolution of 1 x 1 mm. Functional scans used a gradient echo-planar pulse sequence, repetition time (TR) = 1800 ms, echo time (TE) = 23 ms, field of view (FOV) = 240 mm, flip angle = 90°. Images were obtained in 34 axial slices^4^ (no gap, sequential acquisition) of thickness = 3.75 mm, matrix = 64 x 64, resulting in an in-plane voxel resolution of 3.75 x 3.75 mm. The FOV included full cortical coverage and partial cerebellar coverage. For each run we collected 100 functional volumes. Five additional volumes were acquired at the start of each run, but were discarded following acquisition.

### fMRI Data Processing

All fMRI data processing was implemented with custom bash scripting, unless otherwise stated. To improve efficiency of our analysis pipeline, we used GNU Parallel (Tange, 2011) to perform many of the steps described below in parallel for multiple subjects/runs. The fMRI data were processed using functions from FEAT (fMRI Expert Analysis Tool) Version 6.00, part of FSL (FMRIB’s Software Library, www.fmrib.ox.ac.uk/fsl). Preprocessing steps included non-brain removal with BET (Smith, 2002), grand-mean intensity normalisation of the entire 4D dataset by a single multiplicative factor, high pass temporal filtering (Gaussian-weighted least-squares straight line fitting, with sigma = 50.0 s), and spatial smoothing (using a Gaussian kernel of FWHM 6 mm). We also performed motion correction using MCFLIRT (Jenkinson et al., 2002) to mitigate the impact of any potential movement during functional runs. We visually inspected motion correction output; our a-priori threshold for excessive motion between contiguous time points was 2 mm, however no runs exceeded this threshold, therefore no runs were removed due to excessive motion.

Following preprocessing, we used FEAT to estimate activation patterns for each trial using the least-squares-all (LSA) method. For each functional run, we constructed a single GLM that included one regressor for each trial in the run. Each regressor comprised a timeseries modelling the period during which the word for that trial was presented (duration: 2500 ms), convolved with a gamma function (lag: 6 sec, sigma: 3 s) as a model of the haemodynamic response function (HRF). Additionally, we included parameters derived from motion correction as regressors of no interest. We defined 30 contrasts of interest, with each contrast comprising one regressor. This procedure generated four contrasts of parameter estimates (i.e., COPE images) for every unique item; one per trial. Because functional data from different runs are rarely perfectly aligned (due to small head movements between runs), we next aligned all the COPEs from each subject to a common native functional space. We used Advanced Normalization Tools (ANTs; http://stnava.github.io/ANTs/) to rigidly align COPEs from the second, third and fourth runs with the example functional volume (example_func in FSL) from the first fMRI run. We additionally masked the aligned data with a COPE image from the first run, thus removing non-overlapping voxels across runs.

Item-level pattern estimation (described as follows) was performed in the MATLAB environment using functions from the CoSMoMVPA toolbox (Oosterhof et al., 2016). We estimated item-level activity patterns for each unique item by averaging across COPEs from that item’s four respective trials. We then subtracted the mean pattern (i.e., the mean value at each voxel across items) from item-level patterns in each condition separately, in order to remove the activation pattern common to all items in each condition (Walther et al., 2016)^5^.

### Representational Similarity Analysis

#### Searchlight Analyses

We performed RSA using a whole-brain searchlight approach, separately for each subject and condition (aloud or silent reading), in each subject’s native functional space. Searchlights were implemented in the MATLAB environment using functions from the CoSMoMVPA toolbox (Oosterhof et al., 2016). Within each searchlight area (spherical searchlight, radius = 3 voxels), we first generated a neural RDM comprising pairwise Pearson correlation distances (i.e., 1 minus the Pearson correlation) between item-level patterns for all items within that condition. Each neural RDM was vectorized and submitted to a GLM with five regressors, with each regressor comprising one of the five (vectorized) hypothesis models described above. Both the neural data and models were standardised using the *z* transform prior to estimating regression coefficients (Oosterhof et al., 2016). Each searchlight analysis generated five whole-brain statistical maps with a beta (β) value at each voxel; one map was generated for each hypothesis model. β values reflected correspondence between the neural RDM at that voxel (computed from patterns in the corresponding searchlight sphere) and the respective hypothesis model. A separate searchlight analysis was performed for each condition, such that each subject was associated with ten β maps in native functional space.

We next transformed each subject’s searchlight output (i.e., whole-brain β maps) from native functional space to template MNI152 space using ANTs (http://stnava.github.io/ANTs/). We first computed a structural-to-EPI transformation matrix by rigidly aligning participants’ high-resolution structural T1 image to their example functional volume from the first fMRI run. We next computed an MNI152-to-structural transformation matrix using linear affine, and then nonlinear methods (the latter implemented using the “SyN” algorithm; Avants et al., 2008)^6^. We then applied the inverse of the structural-to-EPI and MNI152-to-structural transformation matrices to each subject’s respective searchlight output, re-sliced to 2 mm isotropic resolution. Each subject was therefore associated with ten β maps in MNI152 space. Following this, we masked out the cerebellum (as defined by the Harvard-Oxford cortical atlas supplied by FSL) from all of the spatially normalised β maps; the motivation for this was that our fMRI scans included only partial cerebellar coverage.

#### Group-Level Analyses

For group-level analyses, we computed voxel-wise Bayes factors BF_10_ to evaluate our hypotheses. This is in contrast to conventional null-hypothesis statistical testing (e.g., permutation cluster statistics), which relies on p values to determine whether or not a null hypothesis may be rejected at each voxel. Unlike p values, Bayes factors quantify the strength of evidence in favour of a null or alternative hypothesis. BF_10_ is a ratio that expresses the likelihood of the data under the alternative hypothesis (H_1_) relative to the null hypothesis (H_0_) (Dienes, 2014), and may range between 0-*∞*. Within this framework, incrementally larger values > 1 indicate greater support for the data under H_1_; incrementally lower values < 1 indicate greater support for the data under H_0_. Recently, a growing number of studies have adopted Bayes factors for group-level neuroimaging analyses, particularly in the context of MVPA (Grootswagers, Robinson, & Carlson, 2019; Grootswagers, Robinson, Shatek, et al., 2019; Kaiser et al., 2018; Matheson et al., 2023; Moerel et al., 2022; Proklova et al., 2019; Teichmann et al., 2022). Teichmann et al. (2022) and Matheson et al. (2023) have argued in favour of Bayes factors over p values in MVPA research, primarily because Bayes factors are actually informative as to strength of evidence (as opposed to dichotomous acceptance or rejection of H_1_), and are not susceptible to multiple comparison problems (Dienes, 2016; Teichmann et al., 2022).

We computed Bayes factors using voxel-wise Bayes *t*-tests implemented with functions from the bayesFactor package (Krekelberg, 2022) in the MATLAB environment, using default JZS priors (Rouder et al., 2009). We first tested whether each type of information was decodable in either condition. Here, we performed a right-tailed one-sample Bayes *t*-test at every voxel against the hypothesis that β was greater than zero; this procedure was repeated independently for each condition and type of information. This produced ten statistical maps of BF_10_ values (2 conditions x 5 information types); hereafter, we will refer to these as *within-condition* Bayes maps. Results from these analyses were used to constrain results from contrasts between aloud and silent reading, described as follows.

We next tested the hypothesis that there was a difference in decodability, for each type of information, between conditions. Here, we performed a two-tailed paired-samples Bayes *t*-test at every voxel against the hypothesis that β values differed between aloud and silent reading. This procedure was repeated independently for each type of information, generating five statistical maps of BF_10_ values; hereon, *between-condition* Bayes maps.

As a convention, Bayes factors are often assigned qualitative labels to express the strength of evidence for H_1_, whereby BF_10_ values < 3.0 are considered to provide “weak” or “anecdotal” evidence, values < 10.0 provide “moderate” evidence, and values >= 10.0 are considered to provide “strong” evidence (Dienes, 2014; M. D. Lee & Wagenmakers, 2014). We chose to only consider voxels where there was at least moderate evidence for any hypothesis (that is, decodability greater than 0, or differences between conditions); we feel that reporting weak evidence would not be particularly valuable, and moreover would distract from data points where there was relatively stronger support for our hypotheses. As such, we thresholded all Bayes maps (both within- and between-conditions) at BF_10_ >= 3.0. To establish directionality of our results concerning differences between conditions, we first computed (for each type of information) two average contrast maps: an aloud > silent contrast representing the average searchlight results (across participants) from the aloud condition minus those of the silent condition; vice-versa for the silent > aloud contrast. These average contrast maps were used to mask the thresholded between-condition Bayes maps, thus generating one Bayes map representing aloud > silent, and one representing silent > aloud, for each type of information. Each contrast map was then masked by the within-condition map for its respective minuend condition; this ensured that the contrast maps only included voxels where information was actually decodable in the minuend condition (i.e., it served to exclude any between-condition differences driven entirely by negative values in the subtrahend condition).

We generated interpretable tables of results using FSL’s cluster function to identify clusters of contiguous voxels, separately for each model and between-conditions contrast. We only considered clusters with a minimum spatial extent of 20 voxels, to avoid our results being skewed by noisy voxels. Tables report the spatial location (in MNI coordinates) of the centre of gravity (COG) for each cluster (ascertained using FSL’s *cluster* function), as well as each cluster’s mean (averaged over constituent voxels) and maximum BF_10_ value. Maximum BF_10_ values represent the strongest available evidence for a given hypothesis (e.g., for a difference between conditions) within a cluster, while we consider mean values to reflect evidential strength across the entire cluster. Tables also report anatomical labels for each cluster; these were identified from the Harvard-Oxford cortical and subcortical atlases using FSL’s *atlasquery* function.

## Results

We performed RSA on fMRI data to investigate decodability of information across five types of information—visual, orthographic, phonological, semantic, and articulatory—during aloud and silent word reading. We used a whole-brain searchlight to characterise, at every voxel, correspondence between the neural data and formal hypothesis models representing each type of information, for each condition separately. We also explored differences in decodability between the two conditions. Below, all references to “moderate” or “strong” evidence describe quantitative benchmarking of mean BF_10_ values that were computed for each cluster, based on conventional norms (Dienes, 2014; M. D. Lee & Wagenmakers, 2014).

Both conditions were associated with decodability of visual, orthographic, and phonological information (though to different degrees—see below), while articulatory information was only decodable in the aloud condition. We did not detect semantic information in either condition.

### Visual Information

Results for visual information are shown in Figure 3 and Table 2. We found moderate evidence for visual decodability in both reading conditions. In the left hemisphere, both conditions elicited clusters spanning approximately the same areas of left anterior temporal cortex (including the planum polare, superior and middle temporal gyri, temporal pole) and bilateral inferior frontal cortex (insula and orbitofrontal cortex). Notably, decodability in anterior temporal areas was more extensive in the aloud condition compared to silent. Medial temporal areas were also implicated in both conditions. In the aloud condition visual information was decodable in the right hippocampus and right parahippocampal gyrus; left-hemisphere homologues of these structures (albeit more anterior portions) showed evidence of visual decodability in silent reading.

**Figure 3.**
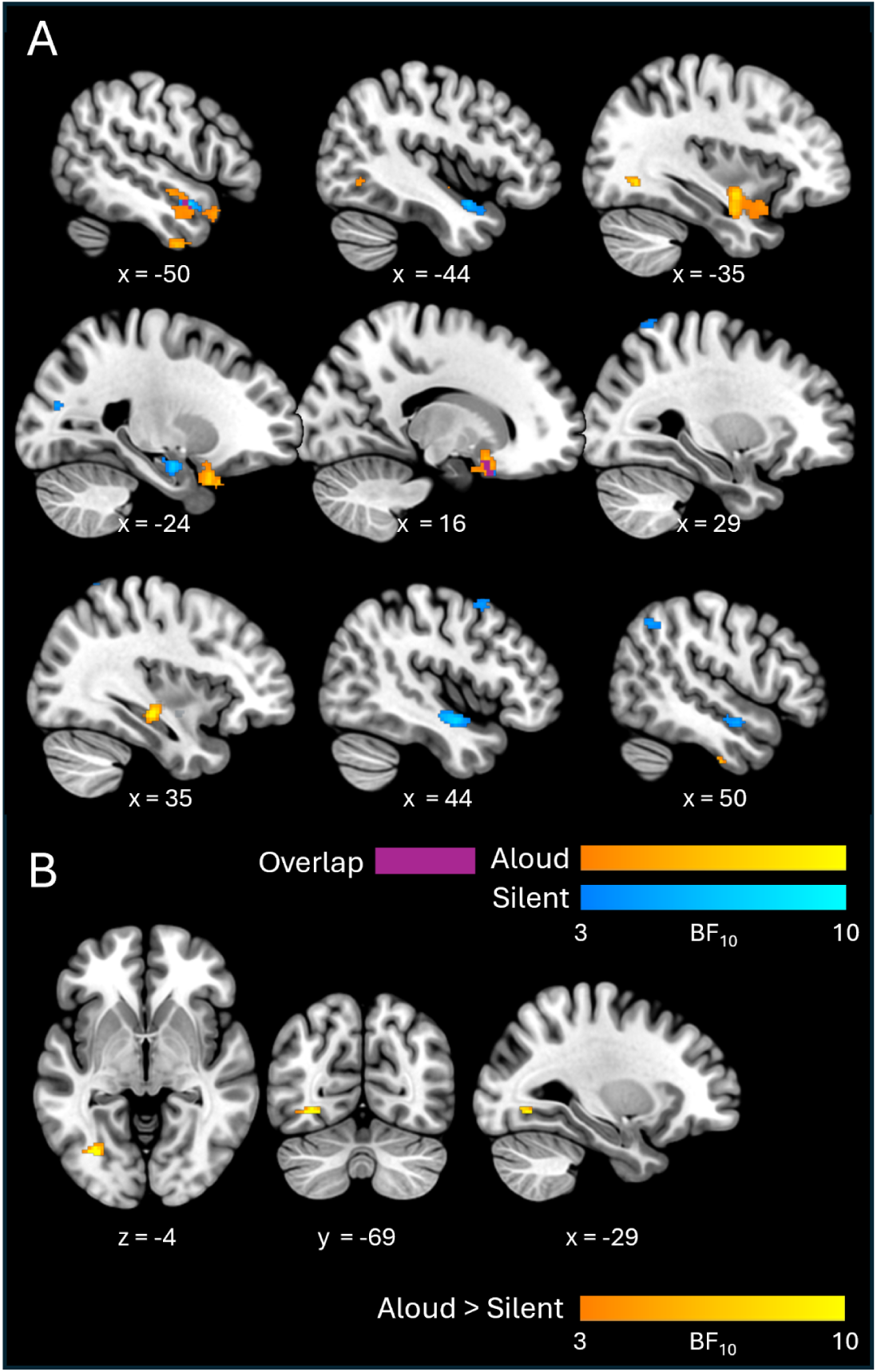
Cross-sectional slices show BF_10_ maps for visual decodability, for individual conditions (A) and between-condition contrasts (B). Axial and coronal views are shown in neurological (left-right) orientation.

Additionally, the aloud condition elicited clusters in ventral aspects of anterior temporal and tempero-occipital cortex—in the anterior portion of the inferior temporal gyrus bilaterally, with another cluster in the posterior portion of the left fusiform gyrus, extending into lateral occipital cortex. By contrast, silent reading was associated with more extensive visual decodability in the right hemisphere. Here, clusters were present in anterior temporal lobe (mirroring effects seen in the left-hemisphere homologue), in the posterior portion of the middle frontal gyrus, and spanning lateral occipital / posterior parietal cortices.

With respect to contrasts between conditions, the aloud > silent contrast revealed a single cluster with moderate evidence (Mean BF_10_ = 8.24) of greater visual decodability in the aloud condition. This cluster was situated in the occipital portion of the left fusiform gyrus (more precisely, on the fusiform bank of the posterior collateral sulcus; Lehman et al., 2016) and extended into lateral occipital cortex. The silent > aloud contrast did not reveal any clusters.

### Orthographic Information

Results for orthographic information are shown in Figure 4 and Table 3. We found evidence for orthographic decodability in both conditions, although decodability was much more apparent (both in terms of quantitative strength of evidence and spatial extent of clusters) in the silent condition.

**Figure 4.**
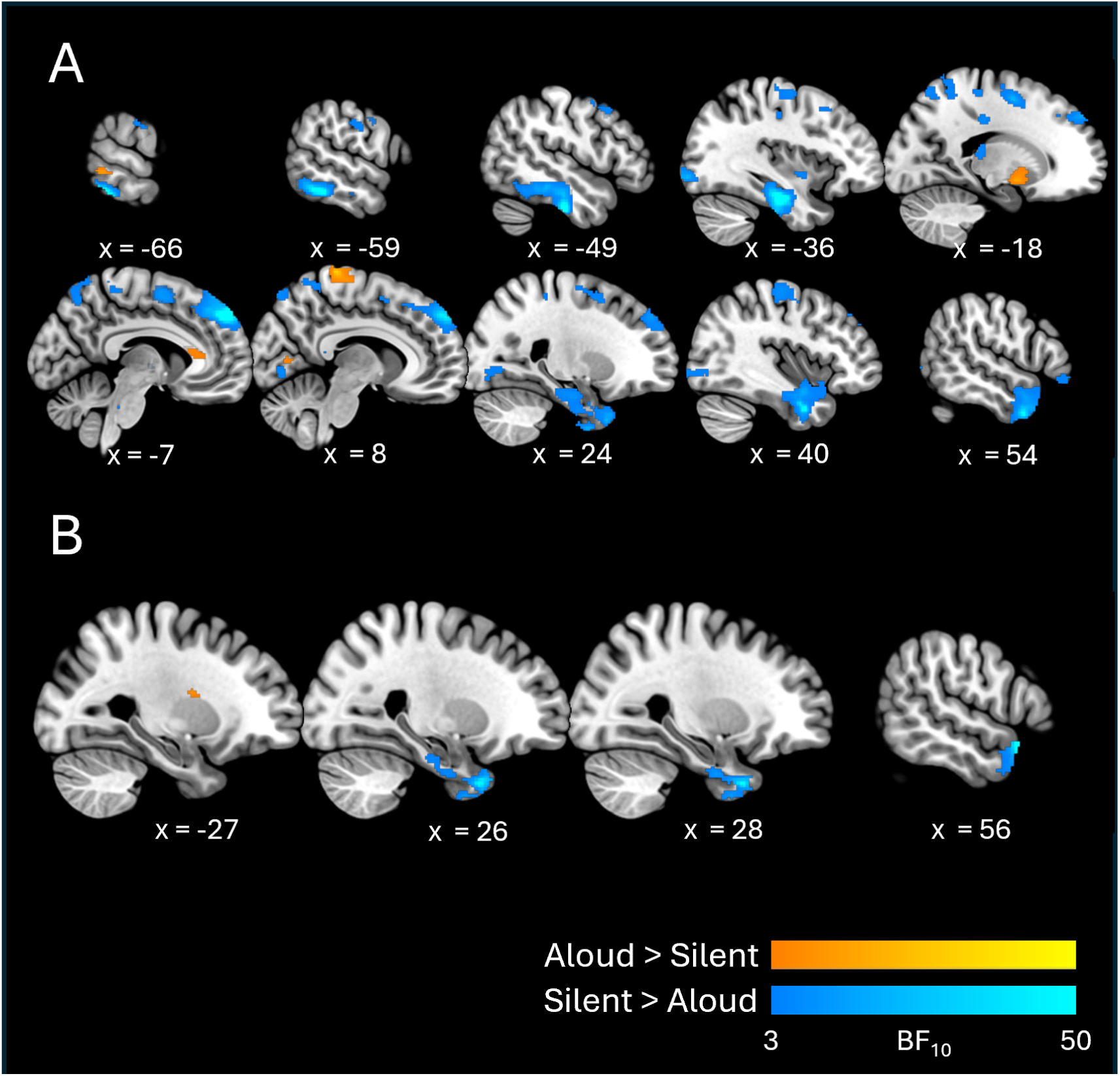
Cross-sectional slices show BF_10_ maps for orthographic decodability, for individual conditions (A) and between-condition contrasts (B).

In the silent condition, the cluster exhibiting the strongest evidence for orthographic decodability (Mean, Max BF_10_ = 22.93, 737) spanned a large area of left inferior and ventral tempero-occipital cortex (including parts of the inferior and middle temporal gyri and the fusiform gyrus), and also extended into left medial temporal areas (parahippocampal gyrus and hippocampus). Another cluster with strong evidence (Mean, Max BF_10_ = 10.59, 263) spanned dorsal frontal cortices bilaterally, including the superior and middle frontal gyri, supplementary motor areas, and primary sensorimotor areas (pre- and postcentral gyri). Besides these, clusters with more moderate evidence (Mean BF_10_ < 10) encompassed broad swathes of medial and lateral occipital cortex bilaterally, medial and lateral parietal cortex bilaterally, right anterior, middle, and medial temporal areas, and right inferior frontal cortex.

By contrast, orthographic decodability in the aloud condition was confined to a few relatively small clusters (with relatively weak evidence: the strongest cluster exhibited Mean, Max BF_10_ = 7.04, 35.5). These clusters were situated mainly in dorsal portions of the pre- and postcentral gyri bilaterally, left lateral tempero-occipital cortex, the lingual gyrus bilaterally, and the anterior portion of the left cingulate gyrus.

With respect to contrasts between conditions, the silent > aloud contrast revealed two clusters with strong-to-moderate evidence (Mean BF_10_ = 12.03, 9.33 respectively) in right anterior temporal lobe. The stronger cluster spanned a relatively large area of anterior ventral temporal cortex, with the peak value (Max BF_10_ = 102) situated at the tip of the temporal pole, extending posteriorly into (anterior portions of) the fusiform and parahippocampal gyri. The second cluster was on the lateral surface, at the border between the temporal pole and superior/middle temporal gyri. The aloud > silent contrast revealed a single cluster with moderate evidence (Mean, Max BF_10_ = 5.73, 10.90) situated in the white matter of the left hemisphere and bordering central opercular cortex (recognising that searchlight spheres may capture activity patterns from patches of grey matter to which they are adjacent; Etzel et al., 2013).

### Phonological Information

Results for phonological information are shown in Figure 5 and Table 4. There was moderate evidence for phonological decodability in both conditions. The aloud condition elicited two clusters: one spanned the left putamen and left insula; the other was situated at the left temperoparietal junction and included the inferior portion of the angular gyrus and the tempero-occipital portion of the middle temporal gyrus. The silent condition also elicited two clusters, one of which was situated in left medial frontal cortex (including mid-cingulate cortex and supplementary motor area) but also extended into the precentral gyrus. The other spanned the middle portion of medial superior frontal gyrus bilaterally.

**Figure 5.**
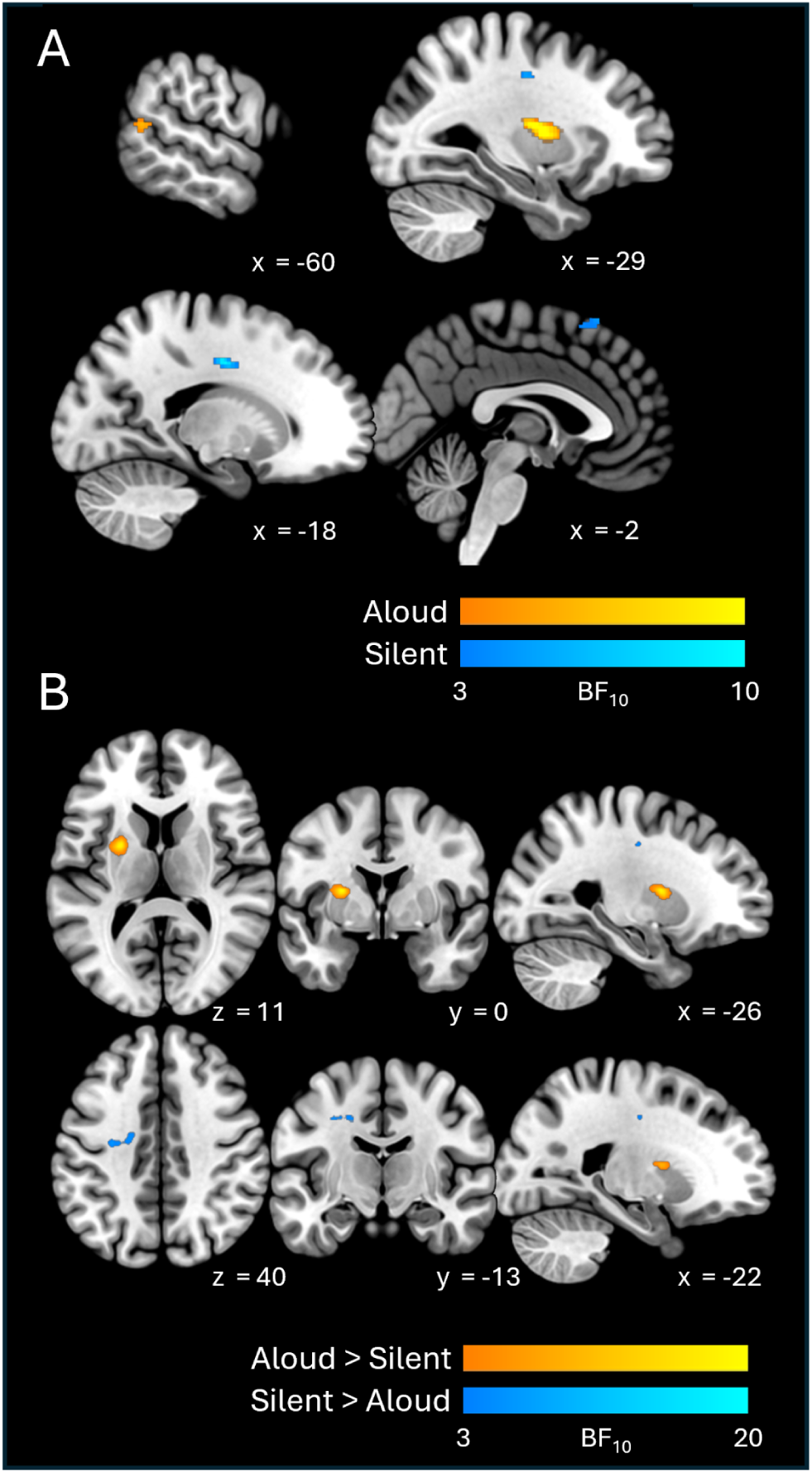
Cross-sectional slices show BF_10_ maps for phonological decodability, for individual conditions (A) and between-condition contrasts (B). Note that axial and coronal views are shown in neurological (left-right) orientation.

**Table 4.**
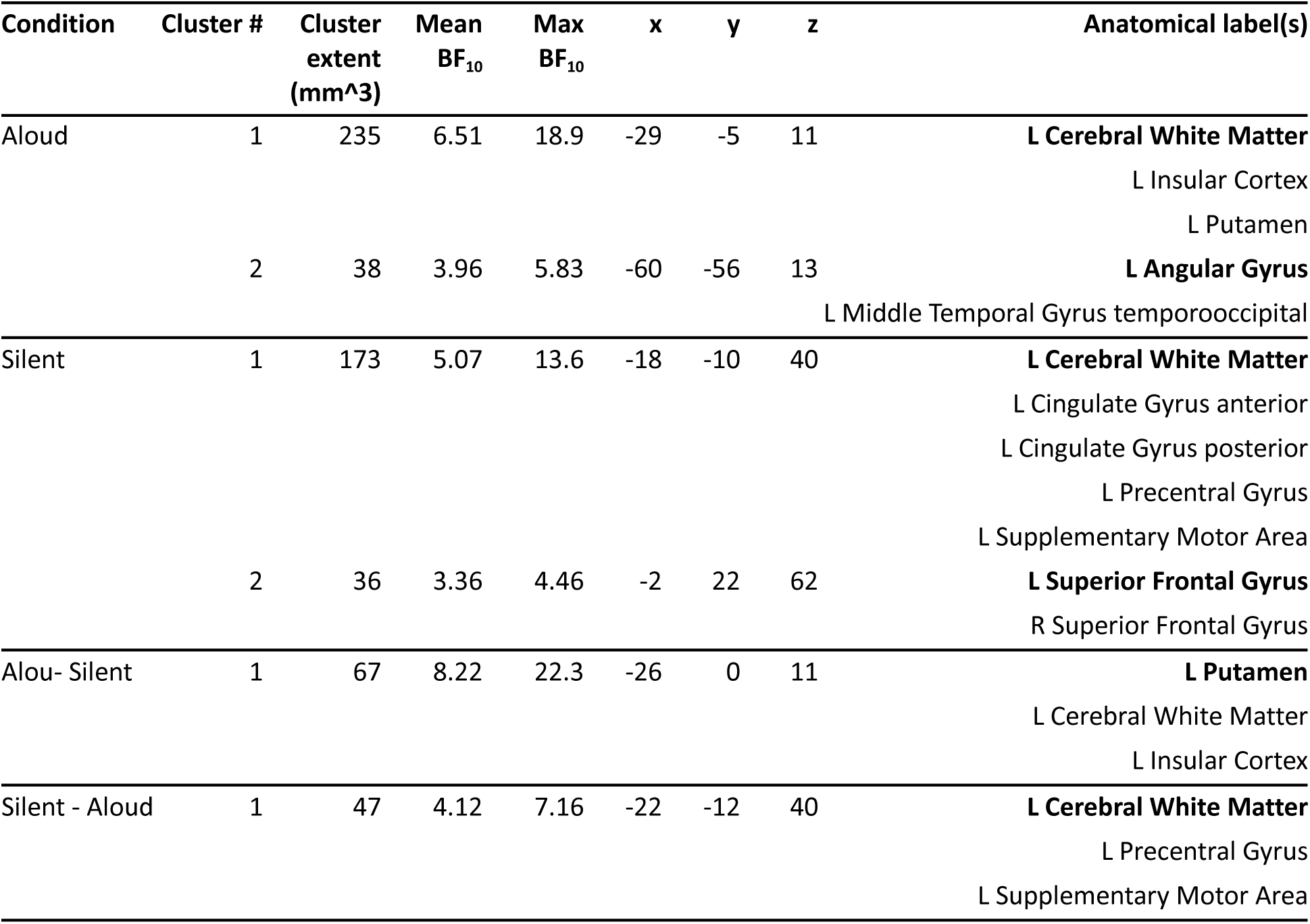
RSA results table detailing clusters for phonological information (i.e., elicited by the phonological hypothesis model). The anatomical labels column shows anatomical structures encompassed by each cluster; entries in boldface indicate the structure in which the cluster’s center of mass (COM) was situated.

With respect to contrasts between conditions, the aloud > silent contrast revealed a single cluster with moderate evidence (Mean, Max BF_10_ = 8.22, 22.30) spanning the left putamen and left insula. The silent > aloud contrast also yielded a single cluster in the left hemisphere, mainly including the left precentral gyrus and underlying white matter.

### Articulatory Information

Results for articulatory information are shown in Figure 6 and Table 5. Both conditions showed moderate evidence of articulatory decodability, however it was much more apparent (in terms of the number of clusters, overall spatial extent, and peak BF_10_ values) in the aloud condition.

**Figure 6.**
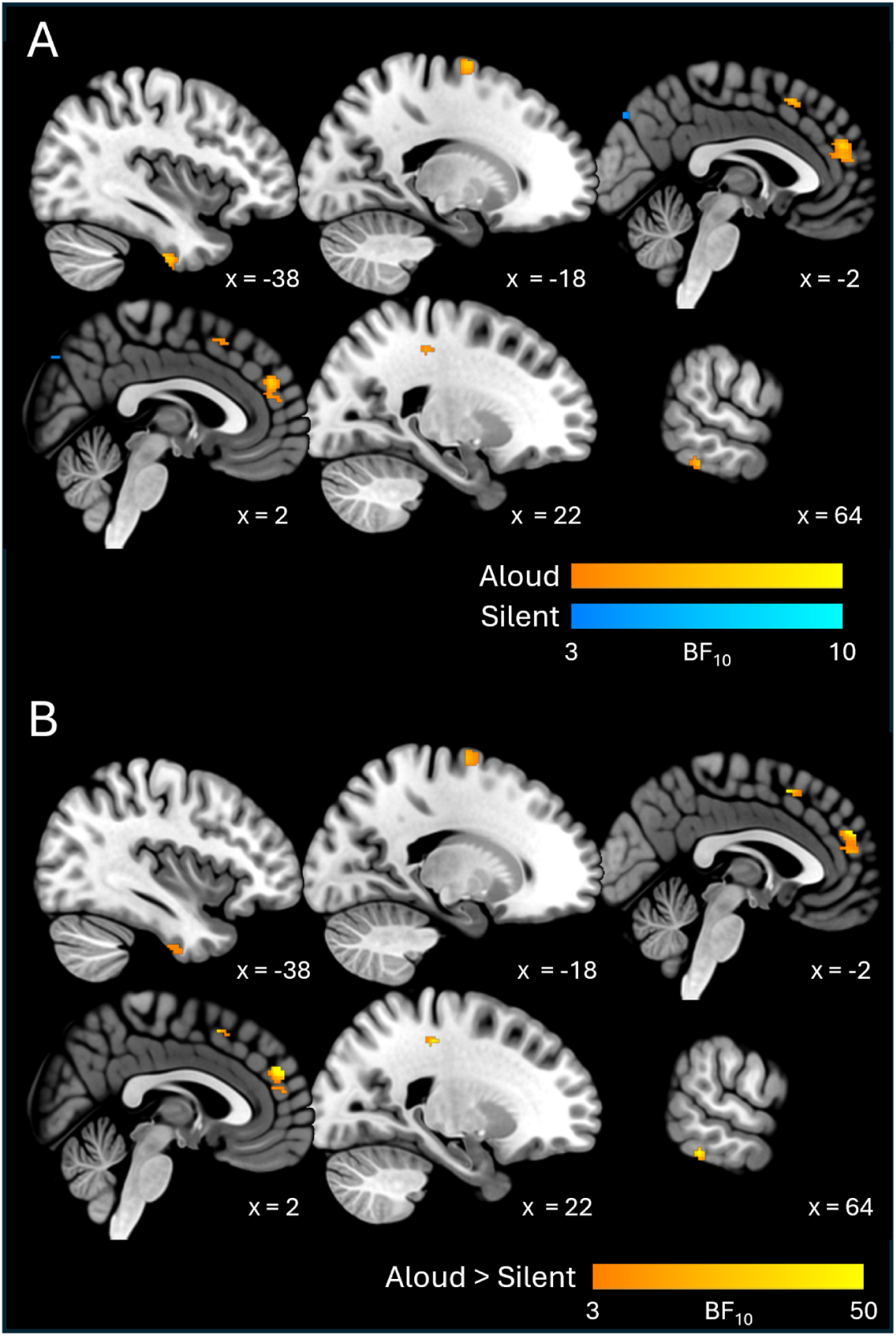
Cross-sectional slices show BF_10_ maps for articulatory decodability, for individual conditions (A) and between-condition contrasts (B).

**Table 5.**
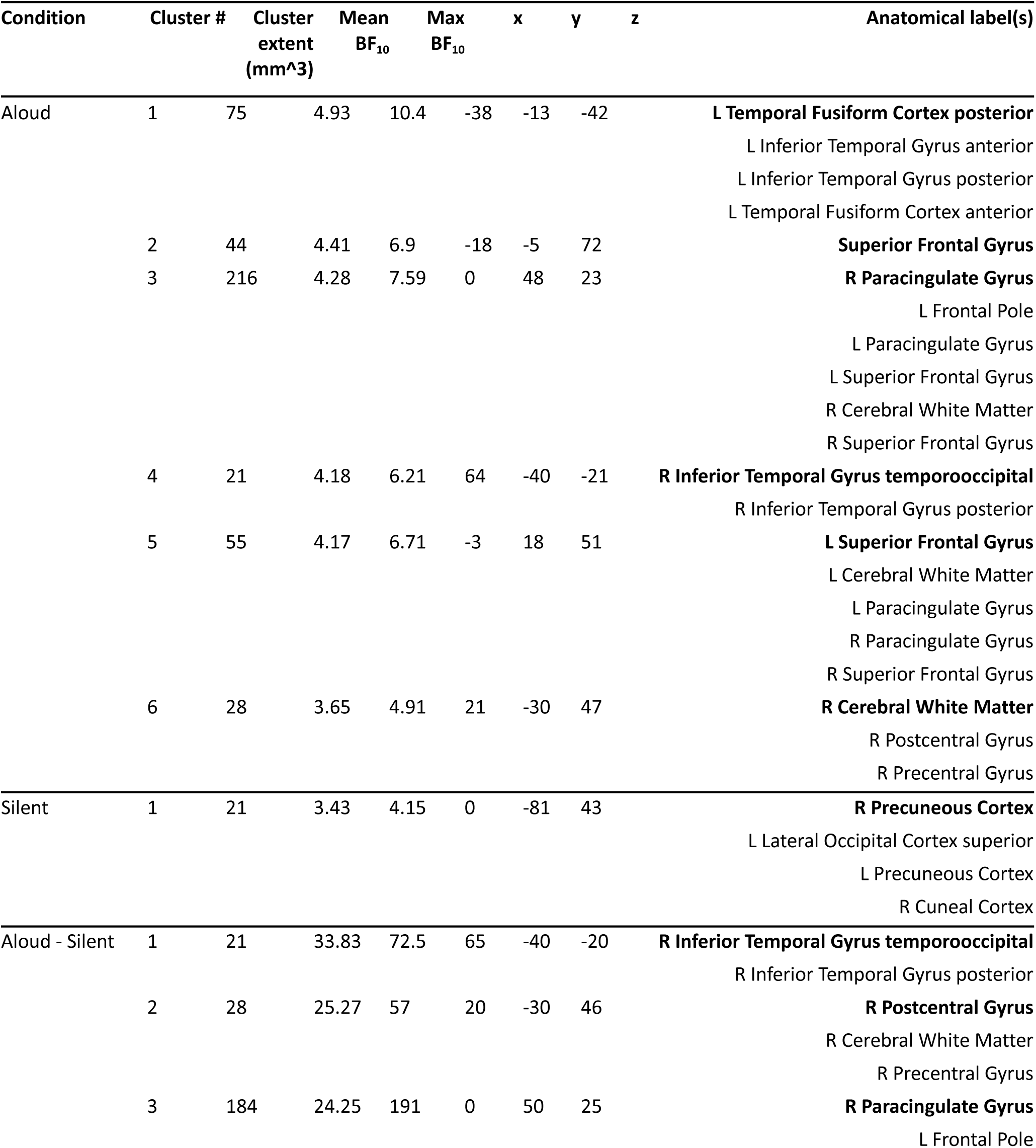
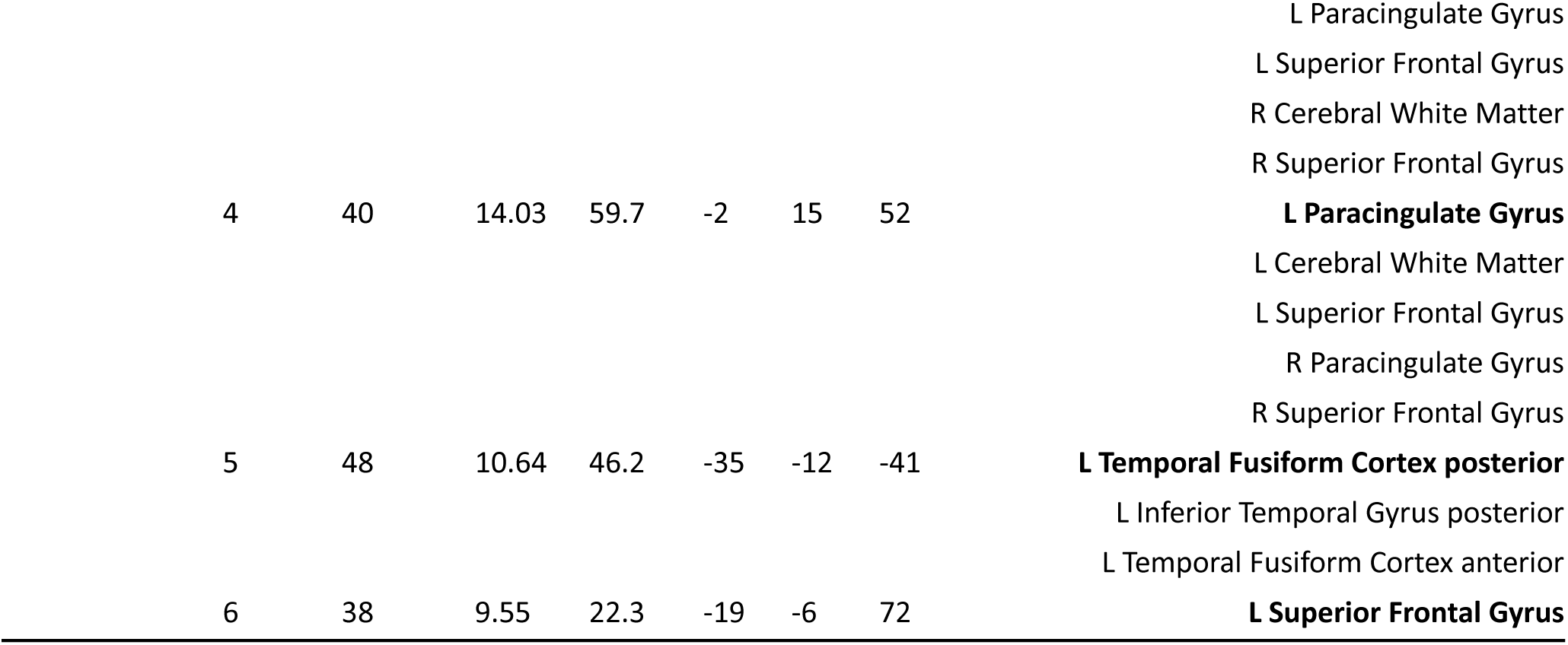
RSA results table detailing clusters for articulatory information (i.e., elicited by the articulatory hypothesis model). The anatomical labels column shows anatomical structures encompassed by each cluster; entries in boldface indicate the structure in which the cluster’s center of mass (COM) was situated.

The aloud condition elicited multiple clusters in the frontal and temporal lobes. On the medial surface of the frontal lobe, two large clusters encompassed areas in both hemispheres. The anteriormost cluster encompassed anterior dorsomedial cortex (superior frontal and paracingulate gyri); the more posterior cluster was situated in the pre-supplementary motor area (pre-SMA). On the lateral surface, one cluster was situated in the posteriormost portion of the superior frontal gyrus (corresponding to the superior portion of dorsal premotor cortex; Schubotz et al., 2010), while the last frontal cluster was situated in the white matter underlying right sensorimotor cortex (again, likely reflecting activity from adjacent grey matter; Etzel et al., 2013). Beyond the frontal lobes, two clusters were situated in the temporal lobes bilaterally: one in the anterior-to-middle portions of the left inferior temporal and fusiform gyri, and another in the tempero-occipital portion of the right inferior temporal gyrus. All of these clusters exhibited moderate evidence. The silent condition elicited a single small cluster (Mean, Max BF_10_ = 3.43, 4.15, spatial extent = 21 voxels) spanning the precuneus bilaterally.

With respect to contrasts between conditions, the aloud > silent contrast revealed clusters that were aligned with all of those elicited by the aloud condition alone. Stated differently: all of the areas showing evidence for articulatory decodability in the aloud condition (described above) also exhibited evidence of increased decodability *relative to* the silent condition. All clusters in the aloud > silent contrast exhibited relatively strong evidence, with the strongest clusters situated in the right hemisphere: in the inferior temporal gyrus (Mean, Max BF_10_ = 33.83, 72.50), primary sensorimotor cortex (Mean, Max BF_10_ = 25.27, 57.00) and anterior medial prefrontal cortex (Mean, Max BF_10_ = 24.25, 191.00)^7^

## Discussion

The current work aimed to characterise differences in the decodability of five types of stimulus-relevant information between aloud and silent single-word reading. We used RSA to compare fMRI data (acquired during each reading task) to formal hypothesis models representing visual, orthographic, phonological, semantic, and articulatory information. Our results revealed differential decodability for all types of information between the two tasks. Interestingly, and contrary to our initial predictions (see Introduction), we did not find evidence that decodability is unilaterally enhanced by aloud reading. Instead, we observed dynamic decodability of information depending on the reading task. Broadly speaking, aloud reading was associated with greater decodability (in terms of the number and spatial extent of clusters as compared to silent reading) of visual, phonological, and articulatory information—this was evident from results for the individual conditions, as well as the aloud > silent contrasts. Silent reading, on the other hand, was associated with greater decodability of orthographic information. We interpret these results to reflect differential weighting of information, depending on cognitive demands imposed by either task.

Below we expand on each set of results and our interpretations therein. Since it is not feasible to address every cluster identified by our analyses, we will focus on data points that we consider to be the most valuable—either those which are directly relevant to our main research objective (identifying differences between aloud and silent reading), or those which are otherwise worthy of comment (e.g., clusters with particularly high strength of evidence and/or spatial extents).

### Visual Information

We found that visual information was decodable in anterior and medial temporal areas, as well as the insula and orbitofrontal cortex, in both reading conditions (albeit more extensively in the aloud condition). This result is surprising, given that these areas are not typically associated with low-level visual processing. On one hand, orthographic information has been detected in anterior temporal lobe by previous RSA studies (Graves et al., 2023; Staples & Graves, 2020; Zhao et al., 2017)—our findings here may reflect early orthographic processes (e.g., recognition of individual characters), which necessarily depend on visual information. Alternatively, this data point might reflect multimodal integration. Many neurobiological models of perception (e.g., Meyer & Damasio, 2009; Patterson et al., 2007; Petrides, 2023) consider anterior temporal lobe to act as multisensory ‘hubs’ where inputs from multiple modalities converge. The insula has similarly been linked to multimodal integration, with a particular emphasis on integrating internal signals (e.g., bodily states) with external sensory input (e.g., Kurth et al., 2010; Salomon et al., 2016). It is possible that our findings in anterior temporal cortex and the insula reflect the presence of visual information as a component of multimodal integration during word reading. The presence of visual information in medial temporal areas may reflect episodic encoding of visual information, consistent with the notion that episodic representations comprise information from multiple modalities (e.g., Jamieson et al., 2016).

Each condition showed evidence of visual decodability in a number of other areas: in ventral temporal and occipital cortices (aloud reading), and in right frontal, temporal, and parietal cortices (silent condition). The aloud clusters in occipital cortex are of particular interest, since this area was also highlighted in the aloud > silent contrast. This contrast revealed a single cluster spanning the posterior portion of the left collateral sulcus and inferior lateral occipital cortex. In the context of reading, these areas have been associated with low-level orthographic analysis (Jobard et al., 2003; Levy et al., 2009); for example, they tend to respond preferentially to consonants versus false fonts (Thesen et al., 2012; Vinckier et al., 2007). We therefore suggest—consistent with the notion that participants allocate more attention to words spoken aloud (Fawcett, 2013; Mama et al., 2018; Ozubko et al., 2012)—this finding reflects greater visual attention to aloud stimuli, which enhances differences between stimuli even at relatively low levels of perceptual processing.

We will not speculate here on the functional significance of the other clusters elicited by each condition. However, the observation that these two conditions elicited clusters in different areas is itself interesting, and speaks to our broader point that informational processing (visual or otherwise) is dynamic and task-dependent.

### Orthographic Information

The orthographic results revealed striking differences between the two reading conditions—while orthographic information was decodable across large patches of frontal, temporal, and parietal cortex in the silent reading condition, the aloud condition revealed only a few small clusters with relatively weak strength of evidence. This suggests that orthographic processing is generally up-regulated for the purposes of silent reading.

As is evident in Figure 4, voxels with the strongest evidence for orthographic decodability in the silent condition were mainly situated in left anterior ventral temporal cortex (with weaker clusters also present in the right-hemisphere homologue), left tempero-occipital cortex, and dorsal frontal cortex bilaterally. The temporal lobe clusters are wholly consistent with previous research. First, left tempero-occipital cortex has long been considered a hub for orthographic processing—for example, the tempero-occipital portion of the left fusiform gyrus houses putative VWFA (e.g., Chen et al., 2019; Martin et al., 2015; Vogel et al., 2012), which is thought to be central to recognition of visual word forms (see Introduction). Previous fMRI-RSA work has also detected orthographic information in this area (Fischer-Baum et al., 2017; Qu et al., 2022). In addition, the more anterior temporal clusters are consistent with previous studies reporting orthographic decodability in middle and anterior ventral temporal cortex bilaterally during single-word reading (Graves et al., 2023; Staples & Graves, 2020; Zhou et al., 2017). The presence of orthographic information in dorsal frontal areas is consistent with previous fMRI-RSA work (e.g., Staples & Graves, 2020 detected orthographic information in the middle frontal gyrus), while some electrocorticography work has shown that the precentral gyrus is sensitive to orthographic information (Kaestner et al., 2021).

The silent > aloud contrast revealed greater orthographic decodability in the right anterior temporal lobe. Notably, evidence for this contrast increased in the posterior-to-anterior direction along the (anterior portion of the) ventral visual pathway (VVP)^8^. This finding is somewhat consistent with Zhao et al. (2017), who identified increasing decodability of orthographic information along the posterior-to-anterior axis of the VVP bilaterally.

As noted earlier, anterior temporal lobe may be considered a hub for multimodal integration. It may be that orthographic information receives greater weighting during multimodal integration in the context of silent reading, perhaps because there are lower demands on phonological / articulatory processing. Another possibility is that silent reading entails relatively greater dependence on print-to-meaning (i.e., direct orthographic-to-semantic) mapping (which has been ascribed to connectivity between the fusiform gyrus and anterior temporal regions; Taylor et al., 2017), as opposed to phonologically mediated access to meaning. While at face value this may seem like a contradictory statement—why would such an effect not be revealed by the semantic analysis?—we reason that extraction of meaning directly from the visual word form should entail sensitivity to orthographic features in the area(s) responsible for that process^9^. Connectionist models of reading consider access to meaning as the sum of parallel inputs from both direct and phonologically-mediated pathways (Harm & Seidenberg, 2004). From this perspective, the ‘division of labour’ between pathways depends on ease of mapping in each^10^. We reason that such division of labour may also be affected by task-related availability of phonological information. That is, if spelling-to-sound computations are already being promoted to meet task demands—as we will argue is the case during aloud reading—then phonologically mediated access to semantics may be assigned greater weighting simply because the relevant computations are already being executed. By extension, reduced demands for phonological information when reading silently may result in *relatively* greater weighting on direct print-to-meaning mapping.

### Phonological Information

Phonological information was detected in both reading conditions, but again, in rather different areas. The aloud condition revealed one cluster spanning the left insula and putamen (also borne out by the aloud > silent contrast), and another at the temporoparietal boundary. Phonological decodability in the insula and inferior parietal cortex is consistent with previous work (Graves et al., 2023; Staples & Graves, 2020), though we are not aware of any RSA studies reporting phonological information in the putamen. This being said, the left putamen has been implicated in speech production (e.g., Oberhuber et al., 2013) and is functionally connected to left-hemisphere cortices associated with language production, comprehension, and reading (see Viñas-Guasch & Wu, 2017 for a meta-analysis). Considering that phonological information is arguably essential for accurate speech production, this first cluster may reflect implementation of articulatory movements necessary to produce the desired sounds. Such processes would be necessary when producing speech, but not (or less so) during silent reading, which may explain why this area was implicated in the aloud > silent contrast.

The silent condition also elicited two clusters, in dorsolateral prefrontal (DLPFC) and medial frontal cortex respectively. The DLPFC cluster is interesting because this area has been implicated in speech planning (Hertrich et al., 2021), but also rule-based inhibitory control (Freund et al., 2021). One possibility is that the presence of phonological information in this area reflects inhibitory phonological “codes”. Considering that participants switched between aloud and silent reading on a trial-to-trial basis, they may have had to actively suppress phonological output on silent trials.

### Semantic Information

Surprisingly, neither condition showed correspondence with the semantic models; this is in contrast to previous research that has identified highly distributed semantic decodability in the context of word reading (e.g., Graves et al., 2023). This null result may have been driven, at least in part, by semantic ambiguity in many of the words in our stimulus list. All of our words were nouns, however some were also verbs (e.g., *address, debate, judge,* etc.). This may have led to inconsistencies in how the words were perceived semantically (and, in turn, inconsistencies in neural representation at the semantic level) across participants, leading to a null effect at the group level. Further work should ensure that stimulus lists are better controlled in terms of syntactic categories.

### Articulatory Information

In the aloud condition, we detected articulatory information in dorsomedial prefrontal cortex (DMPFC) and the pre-supplementary motor area (pre-SMA) bilaterally, left dorsal premotor cortex and anterior ventral temporal lobe, and right postcentral gyrus and inferior tempero-ocipital cortex. These same areas were implicated in the aloud > silent contrast.

DMPFC, pre-SMA, and dorsal premotor cortex are broadly associated with planning and cognitive control of speech processes, amongst other functions (Bourguignon, 2014; Hartwigsen et al., 2013; Hertrich et al., 2021). DMPFC in particular has been linked to domain-general online response monitoring. For example, in the context of a Stroop task, DMPFC may monitor (and resolve) conflicting information arising on a trial-to-trial basis (Freund et al., 2021; C. Kim et al., 2013; A. W. MacDonald et al., 2000). With respect to its role in language production, one study has linked DMPFC to learning arbitrary associations between visually presented stimuli and orofacial and vocal responses, as well as physically performing those responses (Loh et al., 2020). These authors also note that pre-SMA was recruited for learning speech-based vocal responses, but not non-speech vocalisations or orofacial movements, suggesting some degree of specialisation for articulatory processes. Pre-SMA has otherwise been associated with selecting and encoding complex motor sequences, particularly in the context of speech production tasks (e.g., Alario et al., 2006; Tremblay & Gracco, 2009, 2010), with some authors regarding pre-SMA as the “starting mechanism for speech” (e.g., Hartwigsen et al., 2013, p. 580). Premotor cortex, meanwhile, is considered a hub for the articulatory (e.g., phonological-to-motor) components of speech production (e.g., Hickok & Poeppel, 2007), while sequencing of speech motor plans has been ascribed to functional connectivity between dorsal premotor cortex and pre-SMA (Hartwigsen et al., 2013).

We reason that articulatory information in DMPFC, pre-SMA, and dorsal premotor cortex reflects top-down planning and maintenance of vocal responses. To speak each word correctly, participants had to plan and coordinate an appropriate articulatory response comprising a unique sequence of movements of the mouth, jaw, and tongue. The presence of articulatory information in pre-SMA and dorsal premotor cortex likely reflects preparation of this complex motor sequence, while DMPFC may be involved in on-line monitoring to ensure that the produced responses match computed phonological information. The cluster in right postcentral gyrus may reflect sensory input elicited by the articulatory movements themselves. It makes sense that all of these processes should be up-regulated during aloud compared to silent reading. Speech planning and monitoring is only necessary when one’s goal is to read the presented word aloud, therefore it logically follows that articulatory information should be present in cortices governing those processes and—more to the point—to a greater extent in aloud reading (when those processes are required) compared to silent reading (when they are not required).

The functional significance of the inferior and ventral temporal clusters is less clear to us. The cluster in anterior left ventral temporal cortex may reflect multimodal stimulus representations being heavily weighted on articulatory information (and relatively more so during aloud versus silent reading), in line with accounts of the production effect which emphasise the role of articulatory / production features in the episodic memory trace for aloud (but not silent) words (Jamieson et al., 2016; MacLeod et al., 2010).

### General Discussion

Our study provides evidence that aloud and silent reading entail differential decodability of multiple types of stimulus information. Broadly, this finding is consistent with the view of embodied and grounded cognition, whereby different experiences (e.g., reading aloud versus silently) give rise to fundamental changes in how stimuli are perceived and evaluated (Matheson & Barsalou, 2018).

We interpret our results to reflect flexible cognitive states during reading, whereby certain types of information are weighted according to the speakers’ goals. The notion that readers might optimally weigh incoming information to meet task demands is not new, and has even been captured by formal models of word reading (Norris, 2006; Norris & Kinoshita, 2012). In the context of our study, we argue that goal-dependent weighting is primarily driven by the demands of reading aloud.

How is such weighting determined? Consider that some types of information are more relevant or useful for certain tasks; consider also that any reading task will recruit cognitive resources that must be allocated in a manner which best serves the reader’s goals. We suggest that, when reading aloud, phonological and articulatory information are weighted more heavily because those features are useful for planning, execution, and monitoring of speech output. When speech production is not required, the aforementioned types of information are less relevant to one’s goal (reading the word), and so receive less weighting; the result is relatively greater emphasis on orthographic information. For example, de-weighting of spelling-to-sound computations may necessitate relatively emphasis on direct print-to-meaning mapping when reading silently. Overall, our findings provide an adaptive view of information processing, whereby each type of information is weighted according to its task relevance.

A subtly different interpretation, and one which aligns with attentional accounts of the production effect (Fawcett, 2013; Fawcett et al., 2023; Mama et al., 2018; Ozubko et al., 2012), is overall reduced cognitive investment when reading words silently. Silent words might be processed relatively superficially—that is, readers can perceive and extract meaning from visually presented words based on orthographic information alone, without needing to invest further cognitive resources in phonological and articulatory computations. Such superficial processing may apply to all stimulus properties, rather than specific types of information that are directly relevant to production. Indeed, this provides a satisfying explanation for (relatively) reduced decodability of visual information in the silent condition: this information is arguably no more relevant to aloud reading than it is to silent reading, therefore decreased decodability may reflect a global decrease in cognitive investment.

Our findings also have implications for research on the production effect (and, more generally, for research on how words are encoded in memory). The production effect is reliably elicited when memory for aloud versus silent reading is tested, including contexts in which participants are not informed in advance that they will be tested on the studied material (P. A. MacDonald & MacLeod, 1998; Zhou & MacLeod, 2021). Thus while we did not explicitly assess participants’ memory for the words in the present experiment (largely because our stimulus list was relatively short, with each word repeated 4 times), the procedures employed in our experiment (other than stimulus repetition) accurately reflect the encoding conditions that reliably give rise to the production effect. Moreover, our findings align with theoretical accounts of the production effect. One account holds that words read aloud are encoded as highly distinctive memory traces comprising sensorimotor information elicited during articulation (Jamieson et al., 2016; MacLeod et al., 2010). Our finding that reading aloud increased decodability of articulatory information appears to support this position. At the very least, we have shown that articulatory information is *available* for encoding at the neural level, which is a major assumption of distinctiveness accounts.

One possibility we must consider is that the results we observed are epiphenomenal to the production effect. Future research should therefore aim to link decodability (as measured in this study) directly to subjects’ behavioural memory performance; this would provide a more complete understanding of which type(s) of information contribute to encoding. Moreover, future research might consider incorporating formal models of cognition. For example, the MINERVA-2 model of memory has proven able to reproduce the production effect by simulating sensory distinctiveness (Jamieson et al., 2016). In a different vein, some models consider cognitive representations as hierarchical, interconnected structures encompassing all types of stimulus information, as opposed to information being represented in terms of discrete processes (Matheson & Barsalou, 2018; Meyer & Damasio, 2009). Applying such models in the context of RSA may provide a valuable means of testing formal cognitive theories of cognitive and neural representations.

## Data And Code Availability

Code for all analyses reported in this manuscript is publicly available on GitHub [1]; additional materials that are necessary for analyses are stored in an Open Science Framework (OSF) repository [2]. Twelve participants consented to their anonymized data being made publicly available; raw data from those participants are available on the OSF repository [2]. Note that the data reported in this manuscript are from the “quickread” experiment described in both repositories.

[1] https://github.com/lbailey25/Production_Effect_MVPA
[2] https://osf.io/czb26/?view_only=86a66caf1d71484d8ef0293cfa2371df

## Competing Interests Statement

The authors have no competing interests to declare.

## Acknowledgements

We wish to thank the following individuals for their assistance with data collection: Matt Rogers, Laura McMillan, Cindy Hamon-Hill. We also wish to thank Philip Cook for his assistance with spatial normalisation of MVPA output. We also wish to thank three anonymous reviewers for their valuable feedback on earlier versions of this manuscript. This work was supported by grants from the Natural Sciences and Engineering Research Council of Canada (NSERC) to GEB and AJN (Grant numbers: RGPIN-2015-04131, RGPIN-2017-05340). LMB was supported by a Killam Predoctoral scholarship.

## CRediT authorship statement

**Lyam M. Bailey:** Conceptualization, Methodology, Software, Formal analysis, Investigation, Resources, Data curation, Writing - original draft, Visualization, Project administration. **Heath E. Matheson:** Conceptualization, Methodology, Software, Writing - review & editing. **Jonathan M. Fawcett:** Conceptualization, Writing - review & editing. **Glen E. Bodner:** Conceptualization, Funding acquisition, **Aaron J. Newman:** Conceptualization, Methodology, Supervision, Project administration, Funding acquisition, Writing - review & editing.

## Footnotes

1 For example, the vowel sound in *wave, cave, save,* etc. has high spelling-sound consistency in English, because *-ave* usually produces the eɪ vowel. By contrast, the vowel sound in *have* has low consistency because it does not obey this conventional spelling-sound mapping.

2 The searchlight method entails “scanning” a relatively large area (which might be the entire brain or a pre-defined ROI of any size) by parcellating it into a series of smaller searchlight areas, with each centered on a single voxel. An activation pattern for each searchlight area is extracted from its constituent voxels, meaning that one may construct a neural RDM (and, subsequently, compare that RDM to any number of hypothesis models) corresponding to every point in the brain/ROI (Kriegeskorte et al., 2008; Oosterhof et al., 2016).

3 We obtained written permission from *Vecticon* to use the generated voice samples in this research.

4 We added 4 axial slices (total = 38) to the protocol for one participant in order to accommodate their entire cerebral cortex.

5 While mean pattern subtraction is a contentious topic (e.g., Diedrichsen et al., 2011; Garrido et al., 2013), we argue that it is necessary in the case of our study. Shared mean activation has been shown to artificially inflate pairwise correlations between patterns (Walther et al., 2016); this in turn will add noise to correlation-based neural dissimilarity matrices. This has major implications for our study, the purpose of which was to compare decodability (that is, correspondence between neural RDMs and hypothesis models) between aloud and silent reading. Aloud reading reliably elicits more univariate activation than silent reading (Bailey et al., 2021; Dietz et al., 2005; Qu et al., 2022); in turn, item-level patterns for words read aloud likely share more common activation than do patterns for words read silently. As a result, we ought to see relatively greater inflation of pairwise correlations between patterns for words read aloud. Stated differently, failing to control for shared activation (within each condition) would likely result in systematically different degrees of noise contributing to decodability in aloud and silent reading, which would obfuscate any comparisons of decodability between these conditions.

6 To be clear: the structural-to-EPI and MNI152-to-structural transformation matrices were computed using the EPI and structural T1 as reference images respectively. While it is more common to compute these transformations in the opposite direction (i.e., EPI-to-structural and structural-to-MNI152, using structural T1 and MNI152 as reference images), we found that the inverse procedure yielded qualitatively better transformation of searchlight maps—that is, better alignment to the MNI152 template, based on visual inspection of registration output.

7 Notably, the evidence for a difference in decodability between aloud and silent reading was consistently stronger than the evidence for decodability in aloud reading alone. Qualitative inspection of the average searchlight maps indicated that, in each of the areas highlighted by the articulatory analysis, neural RDMs from the silent condition exhibited negative correlations with the articulatory model. This would have resulted in greater numerical differences in data-model correspondence (regression coefficients) between the two conditions, compared to the difference in data-model correspondence from zero in the aloud condition. Why might these negative correlations occur? It may be that the true correspondence between data in the silent condition and the articulatory model was zero, but that noise drove the numerical values below zero. Alternatively, it may be a meaningful effect—for example, activity patterns in the silent condition may be systematically anticorrelated with the model in question (i.e., pairs of words that were more similar in terms of neural activation were more dissimilar in terms of the model, or vice-versa). These considerations are rather beyond the scope of this manuscript. However, we do not feel that either possibility undermines our interpretations of the aloud > silent contrast—whatever the true cause of negative correlations in the silent condition, we can still confidently say that decodability of information was greater in the aloud condition compared to silent.

8 The VVP runs from early visual cortex to anterior ventral temporal cortex, with subsequent projections to prefrontal areas (Kravitz et al., 2013)

9 Importantly, our interpretation does not imply deeper semantic processing in the silent condition (as we did not observe silent > aloud semantic decodability); rather, it concerns the *routes* by which meaning is extracted.

10 For example, sound-meaning mapping is ambiguous in the case of homophones (e.g., ewes-use), and may be resolved by assigning greater weighting to the direct pathway (Harm & Seidenberg, 2004)

## Notes

https://github.com/lbailey25/Production_Effect_MVPA

https://osf.io/czb26/?view_only=86a66caf1d71484d8ef0293cfa2371df

### Competing Interest Statement

The authors have declared no competing interest.

### Summary of Updates

The revised manuscript reports results for individual conditions (version #1 only had results for between-condition contrasts). We have modified our analyses such that phonological information is now captured in terms of acoustic properties. We have revised the Results and Discussion to reflect the new content.

