## Supplementary methods and tables for "Differential weighting of information during aloud and silent reading: Evidence from representational similarity analysis of fMRI data"

### Supplementary Materials

#### Details of dissimilarity measures used for construction of hypothesis models

For our visual measure, we first created an image of each word as it was presented in the experiment using the Pillow package (Umesh, 2012). Each image was then binarized (0 for background pixels, 1 for text pixels) and converted to a vector; we computed visual dissimilarity as the Pearson correlation distance between these vectors (as per Kriegeskorte et al., 2008). For orthography, we took a similar approach to Fischer-Baum et al., (2017): features were unconstrained open character bigrams, computed using the wordkit package (Tulkens et al., 2018), and dissimilarity was calculated as Pearson correlation distance between feature vectors.

For phonology, we used the method described by Bartelds et al. (2020), which is summarized in the manuscript proper. To this end, we collected voice samples from seven volunteers of similar demographic characteristics to our main study sample (three female, two male, one non-binary, age range 20-34)[1]. All volunteers reported that they were born and raised in North America (mostly Canada; one volunteer was from the US) and that English was their dominant language. We reasoned that these volunteers' speech patterns ought to be very similar to those of our fMRI participants, considering that our study took place in Canada. Each volunteer recorded voice samples in a small noise-shielded booth at Dalhousie University. Voice samples were recorded using *Audacity* (version 3.5; https://www.audacityteam.org/) operating on a laptop computer connected to a Logitech YETI USB microphone. Each volunteer was asked to read the full list of study words (S Table 1) aloud in a neutral voice, in alphabetical order. Each volunteer read the full list of words twice, thus providing two samples for every word in the list. In addition to these recordings, we also obtained AI-generated voice samples using the text-to-speech tool provided by the online platform *Vecticon* (https://vecticon.co). We generated speech samples using four of *Vecticon*'s built-in voices: Clara (female, Canadian), Liam (male, Canadian), Jenny (female, American), Davis (male, American). We selected these voices

---

[1] All volunteers gave permission for their voice samples to be used for analysis purposes, and for their anonymized recordings to be made publicly available in our online data repository.

because they all simulate adult speakers with North American accents, and therefore (as with our volunteers) should approximately reflect the speech patterns of our participants.

For semantic models, we used the the gensim package (Řehůřek & Sojka, 2010) to extract vectors from the publicly available glove-wiki-gigaword-300 model (https://github.com/RaRe-Technologies/gensim-data); a pre-trained model of natural language processing based on the Global Vectors for Word Representation algorithm (GLoVe; Pennington et al., 2014). We computed semantic dissimilarity as the cosine distance between vectors, consistent with prior RSA work using so-called word2vec semantic models (e.g. Carota et al., 2021; Guo et al., 2022; Liu et al., 2021; Tong et al., 2022; Wang et al., 2018).

Articulatory models were created using the PhonologicalCorpusTools package (Hall et al., 2019) in conjunction with the Irvine Phonotactic Online Dictionary (IPHOD; Vaden et al., 2009). Each word was transcribed into a string of its constituent phonemes; in turn, each phoneme was represented by a set of features as defined by Hayes (2008). Features reflected actions of the mouth, tongue, and vocal tract required to produce that phoneme (e.g., a phoneme may be voiced or voiceless, has a particular place of articulation, entails some degree of constriction by the lips, teeth, and vocal tract, etc.); thus, we considered them to reflect articulatory components of speech. Articulatory dissimilarity was computed as the edit distance between transcribed phoneme strings (that is, the number of additions, substitutions and/or deletions required to convert one phoneme string to another), weighted according to the number of features shared between constituent phonemes (Fontan et al., 2016). To control for confounding effects of word length, pairwise dissimilarity values were normalized by dividing the computed edit distance by the length of the longest word in the pair (as per Beijering et al., 2008; Schepens et al., 2012).

**Details of measures for extraneous properties**

As described in the manuscript proper, we computed measures of the following uncontrolled properties: imageability, morphological consistency, concreteness, grapheme-to-phoneme consistency, and syntactic category. Numerical imageability and concreteness ratings for each word were obtained from Scott et al. (2019) and Brysbaert et al., (2014) respectively, and dissimilarity between words was computed as the absolute numerical difference between words in both cases. For morphological complexity, we counted the number of morphemes in each word; dissimilarity was computed as the absolute difference in morpheme counts. For grapheme-to-phoneme consistency, words were vectorised according to their syllable-wise spelling-to-sound consistency (Chee et al., 2020) and dissimilarity was computed as the Euclidean distance between vectors (we found that using Euclidean distances produced overall more normally distributed hypothesis models, compared with other metrics such as correlation or cosine distance). For syntactic category, each word was assigned a binary value: 0 if the word was a noun only, 1 if the word was both a noun and a verb. Dissimilarity was again computed as the absolute difference in value.

#### Supplementary tables

*S Table 1.* List of words used in the current study

| | | | | | |
|---|---|---|---|---|---|
| account | century | garden | language | pebble | summer |
| answer | department | handle | machine | powder | ticket |
| author | education | industry | message | record | turnip |
| beauty | envelope | journey | ocean | sailor | valley |
| campaign | forest | kingdom | painting | speech | wheat |

*S Table 2*. Values show correlations between hypothesis models (used for RSA) and potentially confounding stimulus properties. Parenthetical values are Bayes factors indicating strength of evidence for each correlation. Abbreviations: G2P consistency = grapheme-to-phoneme consistency; Morph. complexity = morphological complexity.

| | **Visual** | **Orthographic** | **Phonological** | **Semantic** | **Articulatory** |
|---|---|---|---|---|---|
| **Concreteness** | 0.06 (0.09) | -0.01 (0.04) | 0.11 (0.61) | 0.15 (6.03) | 0.03 (0.05) |
| **G2P consistency** | -0.04 (0.06) | -0.03 (0.05) | 0.22 (2072.86) | -0.14 (3.38) | 0.12 (0.79) |
| **Imageability** | -0.02 (0.06) | -0.06 (0.07) | 0.21 (7.35) | 0.15 (0.82) | 0.11 (0.21) |
| **Morph. complexity** | 0.09 (0.22) | < 0.01 (0.04) | 0.14 (3.43) | 0.02 (0.04) | -0.05 (0.07) |
| **Syntactic category** | -0.04 (0.06) | -0.07 (0.11) | -0.04 (0.05) | < -0.01 (0.04) | -0.06 (0.08) |
